# Ezh2-dependent epigenetic reprogramming controls a developmental switch between modes of gastric neuromuscular regulation

**DOI:** 10.1101/486423

**Authors:** Sabriya A. Syed, Yujiro Hayashi, Jeong-Heon Lee, Huihuang Yan, Andrea Lorincz, Peter R. Strege, Gabriella B. Gajdos, Srdjan Milosavljevic, Jinfu Nie, Jüri J. Rumessen, Simon J. Gibbons, Viktor J. Horvath, Michael R. Bardsley, Doug D. Redelman, Sabine Klein, Dieter Saur, Gianrico Farrugia, Zhiguo Zhang, Raul A. Urrutia, Tamas Ordog

## Abstract

Physiological interconversion between specialized cell types has only been described in a few mammalian tissues and the mechanisms remain obscure. Using genetic lineage tracing during postnatal development and *in-vitro* models we demonstrate conversion of gastric interstitial cells of Cajal (ICC), regulatory cells that electrically pace phasic contractions and mediate nitrergic and cholinergic neural control of smooth muscle cells, into phenotypically distinct “fibroblast-like” interstitial cells (FLC), which only mediate purinergic signaling. Mechanistically, we find this transition to be epigenetically governed by H3K27 trimethylation of cell identity-related promoters whose susceptibility to repression is predicted by H3K27 acetylation patterns in ICC. The phenotypic switch was reversible by inhibition, knockdown or *in-vivo* genomic inactivation of the polycomb H3K27 methyl-transferase Ezh2. These results demonstrate a role for Ezh2-mediated epigenetic repression in physiological mammalian transdifferentiation and identify FLC as a reserve from which ICC can potentially be restored in common gastrointestinal disorders where ICC are depleted.

**GRAPHICAL ABSTRACT:** 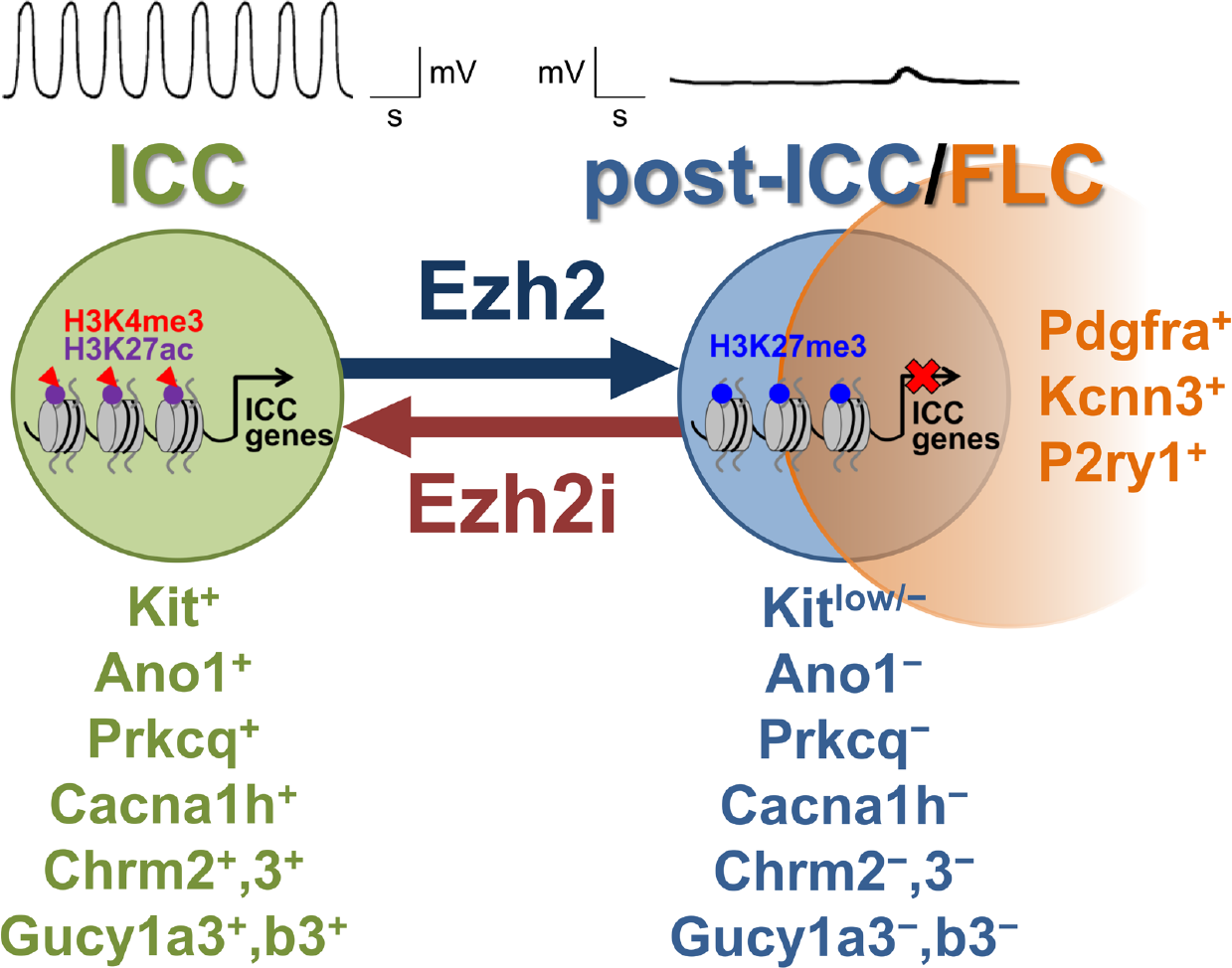

**HIGHLIGHTS:**

- Gastric pacemaker cells (ICC) transdifferentiate into quiescent cells (FLC) in vivo
- ICC-to-FLC shift switches neural control from nitrergic/cholinergic to purinergic
- Ezh2-mediated H3K27me3 represses cell-identity genes during ICC-to-FLC transition
- Ezh2 inhibition restores ICC numbers, phenotype and function

**eTOC BLURB:** Syed et al. find aging to cause transdifferentiation of gastric pacemaker cells (interstitial cells of Cajal, ICC), which also communicate cholinergic and nitrergic neurotransmission to smooth muscle cells, into quiescent “fibroblast-like cells” (FLC), which only mediate purinergic signals. This switch is governed by Ezh2, whose inhibition can reverse ICC depletion.

## INTRODUCTION

Transdifferentiation is a form of cellular plasticity that involves interconversion between specialized cell types with or without intervening de-differentiation into a primitive stem-like state (Merrell and Stanger, 2016). Homeostatic transdifferentiation occurring in response to developmental conditions or injury has been demonstrated in invertebrates and amphibians but only few examples have been reported in mammalian tissues (Chera et al., 2014; Yanger et al., 2013). Furthermore, the central question of how alterations in chromatin structure set up and maintain heritable transcriptional programs underlying mammalian transdifferentiation remains unanswered.

We investigated this problem in the gastrointestinal (GI) neuromuscular apparatus, where transdifferentiation of interstitial cells of Cajal (ICC) into smooth muscle-like cells (Chang et al., 2001; Mei et al., 2009; Torihashi et al., 1999) has been proposed to contribute to the ICC depletion occurring in virtually every major GI neuromuscular disorder including inflammatory diseases, diabetic gastroparesis, aging-related dysfunction (Farrugia, 2008; Ordog, 2012) and “functional” GI disorders (Angeli et al., 2015), which lack other causative lesions. ICC pace all peristaltic and non-propulsive smooth muscle contractile activity by generating electrical slow waves via the calcium-activated chloride channel Ano1 and coupled mechanisms (Huizinga et al., 2014; Klein et al., 2013; Sanders et al., 2014). In addition, ICC that intercalate between smooth muscle cells and enteric nerve fibers play dominant roles in mediating cholinergic excitatory and nitrergic inhibitory regulation of smooth muscle function by the enteric and systemic autonomic nervous systems (Groneberg et al., 2015; Klein et al., 2013; Lies et al., 2014; Sanders et al., 2014). Transcriptional programs underlying ICC phenotypes and functions are orchestrated by signaling from the receptor tyrosine kinase (RTK) Kit, which, in conjunction with the ets family transcription factor Etv1, plays a permissive role in ICC differentiation (Chi et al., 2010; Torihashi et al., 1997) from embryonic and adult precursors (ICC stem cells; ICC-SC) (Bardsley et al., 2010; Lorincz et al., 2008).

Despite their high prevalence, GI neuromuscular disorders lack effective, mechanism-based, curative therapy (Ordog, 2012) and cost billions of dollars in health care spending each year (2009). ICC loss is central to the pathogenesis of many of these disorders (Farrugia, 2008; Ordog, 2012). Therefore, the possibility of transdifferentiation, rather than cell death, underlying the ICC loss observed in these diseases raises hope for new mechanistic, cell-based therapeutic options. Although in various experimental models ICC-like cells with aberrant phenotypes have been reported (Chang et al., 2001; Faussone-Pellegrini et al., 2006; Lorincz et al., 2008; Malysz et al., 1996; Mei et al., 2009; Torihashi et al., 1999), conclusive evidence of long-term ICC survival following loss of Kit and ICC transdifferentiation has been elusive.

Our goal in this study was to positively identify ICC fates and determine their epigenetic mechanisms and reversibility during early postnatal development in mice. Since expression of Kit and the closely related RTK Pdgfra are inversely correlated during ICC differentiation (Bardsley et al., 2010; Hayashi et al., 2015; Kurahashi et al., 2008), we hypothesized that following loss of Kit, ICC may again upregulate Pdgfra expression and transdifferentiate toward Pdgfra^+^ interstitial cells (“fibroblast-like cells”, FLC) rather than smooth muscle cells. FLC have a less diverse functional repertoire compared to ICC. FLC do not generate electrical slow waves and express several purinergic receptors and the Kcnn3 (also known as SK3) intermediate/small conductance calcium-activated potassium channel (Iino et al., 2009; Peri et al., 2013), which is thought to carry the FLC-specific apamin-sensitive currents underlying purinergic inhibitory neuromuscular neurotransmission in GI muscles (Sanders et al., 2014). Furthermore, unlike ICC, FLC may not be reduced in neuromuscular diseases (Grover et al., 2012). Thus, ICC and FLC are different, highly specialized cell types mediating distinct modes of enteric neuromuscular control and responding differently to stress.

Using genetic lineage tracing and novel cell models, here we demonstrate that during the first 3-5 months of life, nearly one-third of ICC lose their phenotype and function with 56% of these “post-functional ICC” (post-ICC) acquiring Pdgfra expression and the more limited functional repertoire of FLC. Integrated transcriptome and epigenome profiling revealed that genes related to ICC identity and function bearing specific patterns of histone 3 lysine 27 acetylation (H3K27ac) became preferentially silenced during the transition of ICC to FLC. We found this transcriptional repression to be mainly mediated by the polycomb repressive complex 2 (PRC2) H3K27 methyltransferase Ezh2. RNA interference-mediated knockdown, pharmacological inhibition and *in-vivo* genomic deletion of *Ezh2* permitted the recovery of functional, Kit^+^ ICC even after the phenotypic transition had taken place. These results demonstrate a previously unrecognized role for Ezh2-mediated epigenetic repression in physiological mammalian transdifferentiation and identify Pdgfra^+^ FLC as a reserve cell population from which Kit^+^ ICC can potentially be restored by epigenetic reprogramming in common GI disorders where ICC are depleted.

## RESULTS

### Lineage tracing reveals ICC survival following loss of Kit expression

To determine the fate of ICC during mouse postnatal development, we performed genetic lineage tracing combined with flow cytometry based on previously established and validated approaches (Bardsley et al., 2010; Chen et al., 2007; Klein et al., 2013; Lorincz et al., 2008) (see **Supplemental Experimental Procedures** for experimental details and reagents). In *Kit*^*CreERT2*/+^*;R26*^*mT-mG/mT-mG*^ mice, Kit-transcribing cells were indelibly pulse-labeled with membrane-targeted green fluorescent protein (mG) expressed in response to Cre-mediated recombination induced by 3-day tamoxifen treatment between postnatal days (P) 8-10. The number and distribution of mG^+^ cells among cells lacking hematopoietic markers (HP^−^; to exclude Kit^+^ mast cells) and expressing combinations of Kit, Cd34 and Pdgfra on their surface was analyzed by flow cytometry in the whole gastric corpus+antrum *tunica muscularis* (**Figure S1A,B**) on P11 (n=7) and P107-160 (adult group; n=5). Between these ages, average counts of ICC, identified by immunolabeling as HP^−^Kit^+^Cd34^−^ cells declined by 57% and Kit immunofluorescence of ICC was reduced (**Figure 1A,B**). Similarly to progeric *klotho* mice (Izbeki et al., 2010), ICC precursors (pre-ICC) including HP^−^Kit^low^Cd34^+^ ICC-SC and HP^−^ Kit^+^Cd34^+^ immature ICC were also reduced (**Figure 1B**). These results indicate a dramatic decline in cells of the ICC lineage occurring during the first 3-5 postnatal months and establish an *in-vivo* model to study ICC fates.

**Figure 1.**
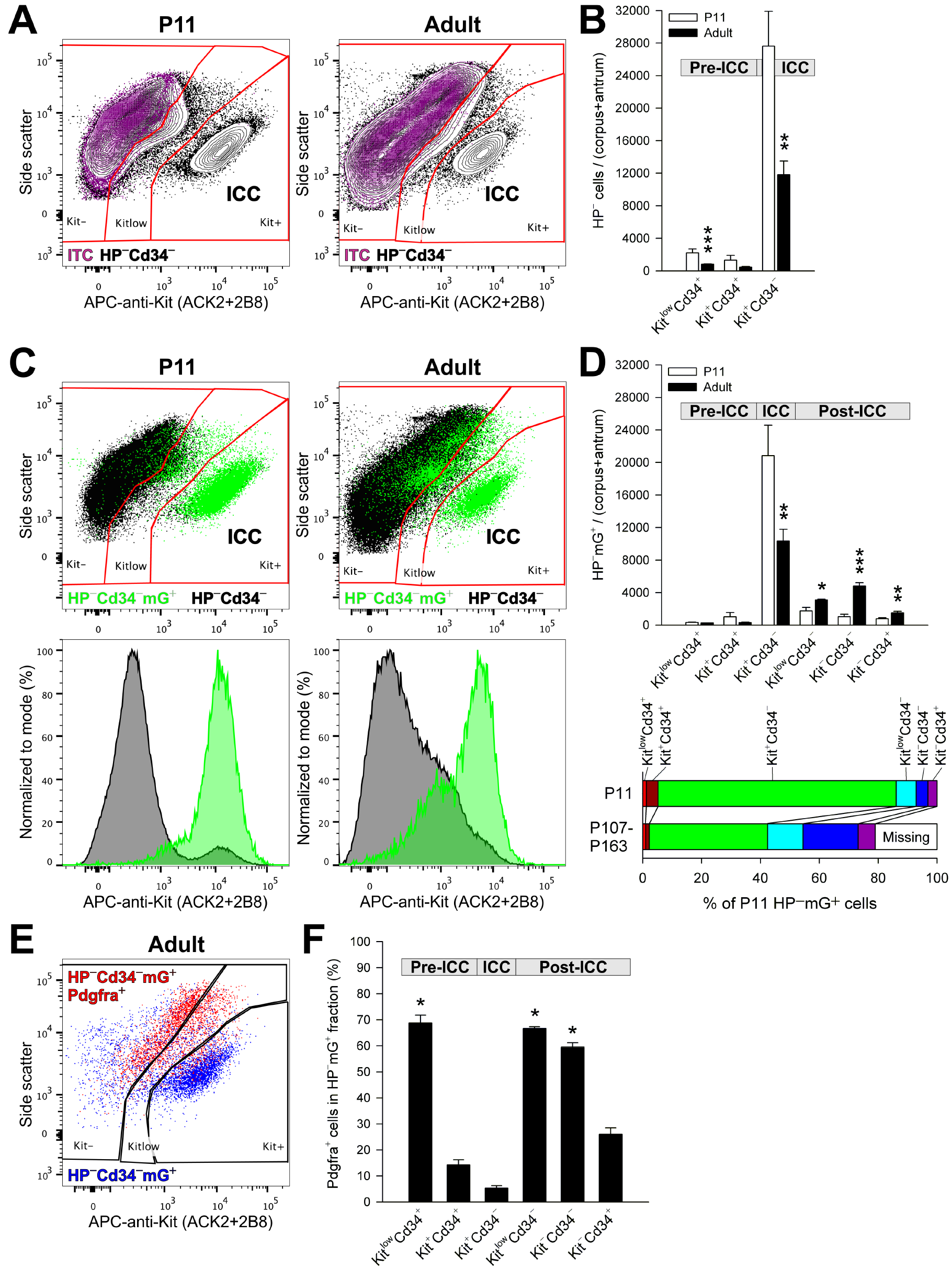
Kit^+^ ICC transition into Kit^low/−^Pdgfra^+^ cells *in vivo*. Panels display representative and quantitative flow cytometry data from the HP^−^ fraction (Cd45 (Ptprc)^−^, Cd11b (Itgam)^−^, F4/80 (Adgre1)^−^ cells) of the gastric corpus+antrum *tunica muscularis* of 7 P11 and 5 adult, tamoxifen-treated *Kit*^*CreERT2*/+^*;R26*^*mT-mG/mT-mG*^ mice. Bar graphs show means±s.e.m.; *P* values are from rank sum tests. (**A**) Representative 2% quantile contour maps showing age-related reduction in the proportion and Kit immunoreactivity of Kit^+^Cd34^−^ ICC. Purple color: cells labeled with an antibody panel where the anti-Kit antibodies were replaced with fluorochrome-matched isotype control antibodies (ITC). (**B**) Age-related reduction of Kit^low^Cd34^+^ ICC-SC (***, *P*=0.003), Kit^+^Cd34^+^ immature ICC (*P*>0.05) and Kit^+^Cd34^−^ ICC (**, *P*=0.01). (**C**) Representative dot plots (100,000 HP^−^Cd34^−^ cells) and histograms showing age-related shift of genetically traced Kit-transcribing cells (mG^+^, green) from Kit^+^Cd34^−^ ICC to Kit^low/−^Cd34^−^ populations. (**D**) Top: Age-related decrease in mG^+^ cell numbers in Kit^low^Cd34^+^ ICC-SC (*P*>0.05), Kit^+^Cd34^+^ immature ICC (*P*>0.05) and Kit^+^Cd34^−^ ICC (**, *P*=0.03) and concomitant increase in mG^+^ counts in the post-ICC populations Kit^low^Cd34^−^ (*, *P*=0.048), Kit^−^Cd34^−^ (***, *P*=0.003) and Kit^−^Cd34^+^ (**, *P*=0.018). Bottom: Age-associated reduction in ICC and pre-ICC and expansion of post-ICC. Data are percentages of average total HP^−^mG^+^ cell count at P11. (**E**) Representative dot plot (7,100 HP^−^Cd34^−^mG^+^ cells) demonstrating inverse relationship between Kit and Pdgfra expression. (**F**) Pdgfra^+^ cell frequencies in mG^+^ cells of the ICC lineage in adults. Similarly to ICC-SC, the majority of Kit^low/−^ post-ICC express Pdgfra. *, *P*<0.05 vs. Kit^+^Cd34^−^ ICC (Dunn’s multiple comparison following Kruskal-Wallis *H* test, *P*<0.001, *H*(5)=26.894). See also **Figure S1**.

At P11 (n=7), 73±4% of HP^−^Kit^+^Cd34^−^ ICC, 70±6% of HP^−^Kit^+^Cd34^+^ immature ICC and 16±2% of HP^−^Kit^low^Cd34^+^ ICC-SC were mG^+^ indicating strong relationship between recombination and *Kit* transcription (see detailed validation of the lineage tracing method in **Supplemental Experimental Procedures**). In adult mice (n=5), the decline in the mG^+^ ICC and immature ICC mirrored their loss determined by immunolabeling (**Figure 1A-D**), with lineage-traced ICC decreasing by 50%. These results demonstrate the high fidelity of our lineage tracing approach.

In adult mice, the decrease in lineage-traced ICC and pre-ICC was associated with significant increases in mG^+^ cells in the HP^−^Kit^low^Cd34^−^ and HP^−^Kit^−^Cd34^−^ fractions, which had been proposed to contain ICC with reduced or lost Kit expression and function (Lorincz et al., 2008), and also in HP^−^Kit^−^Cd34^+^ cells containing a small subset of FLC (Iino and Nojyo, 2009) (**Figure 1C,D**). The net age-related gain in these cells, collectively termed as post-ICC, accounted for 56% of the mG^+^ ICC loss and represented 28% of all mG^+^ ICC detected at P11.

In all, 79% of the mG^+^ cells detected at P11 were accounted for in the adults (**Figure 1D**, lower panel). These results indicate that in mice during the first 3-5 months of postnatal life, 21% of cells of the ICC lineage likely die, about half survive as ICC and the remaining 28% survive as Kit^low/−^ cells distinct from ICC and pre-ICC.

### Lineage tracing reveals ICC phenotypic transition into Pdgfra^+^ cells

Since Pdgfra is considered to be the key RTK and a well-established marker of FLC (Iino et al., 2009; Sanders et al., 2014), we investigated Pdgfra expression in genetically traced post-ICC. Consistent with our previous reports, in adult mice, Pdgfra and Kit expression were inversely correlated in the HP^−^Cd34^−^mG^+^ population (**Figure 1E**) and in the mG^+^ subsets of pre-ICC and ICC (**Figure 1F**). In mG^+^ post-ICC, Pdgfra^+^ cell frequencies were 67±1% (Kit^low^Cd34^−^), 59±2% (Kit^−^Cd34^−^) and 26±2% (Kit^−^Cd34^+^) (**Figure 1F**). These results demonstrate that during the first 3-5 months of postnatal life, more than half of ICC that survive following the downregulation of Kit expression acquire the key FLC marker Pdgfra.

### Conditionally immortalized ICC retain key ICC traits

Functional, Kit^+^ ICC cannot be maintained and propagated in primary cultures beyond a few days even in the presence of Kit ligand (Kitl)-expressing feeder cells (Ordog et al., 2004; Rich et al., 2003; Ward et al., 2000). Therefore, to replicate the ICC-to-post-ICC transition *in vitro* and facilitate mechanistic studies, we established two clonal, conditionally immortalized ICC cell lines by fluorescence-activated cell sorting (FACS) (ICL2A and ICL2B; **Figure 2A, Supplemental Experimental Procedures**) and a Kit^+^ ICC-dominated cell line (ICL1) from a homozygous, P116 Immortomouse by immunomagnetic selection (MACS). In early passages (<8), all lines displayed key ICC characteristics including dendritic morphology and formation of interconnected networks via slender processes (**Figure 2B**), expression of Kit protein and mRNA (**Figure 2C,D**), as well as ultrastructural features considered hallmarks of ICC (Sanders et al., 2014) (**Figure 2E, Figure S2A, Table S1**). By flow cytometry, immunocytochemistry and Western blotting performed between passages 11-16 the clonal ICC lines also expressed the ICC marker Cd44 (Chen et al., 2007; Lorincz et al., 2008) but did not contain smooth muscle myosin heavy chain (Myh11)- expressing smooth muscle cells, Cd34^+^ cells including ICC-SC, Cd45 (Ptprc)-expressing hematopoietic cells, glial fibrillary acidic protein (Gfap)-expressing glial cells or protein gene product 9.5 (Pgp 9.5; Uchl1)-expressing neurons (**Figure S2B**).

**Figure 2.**
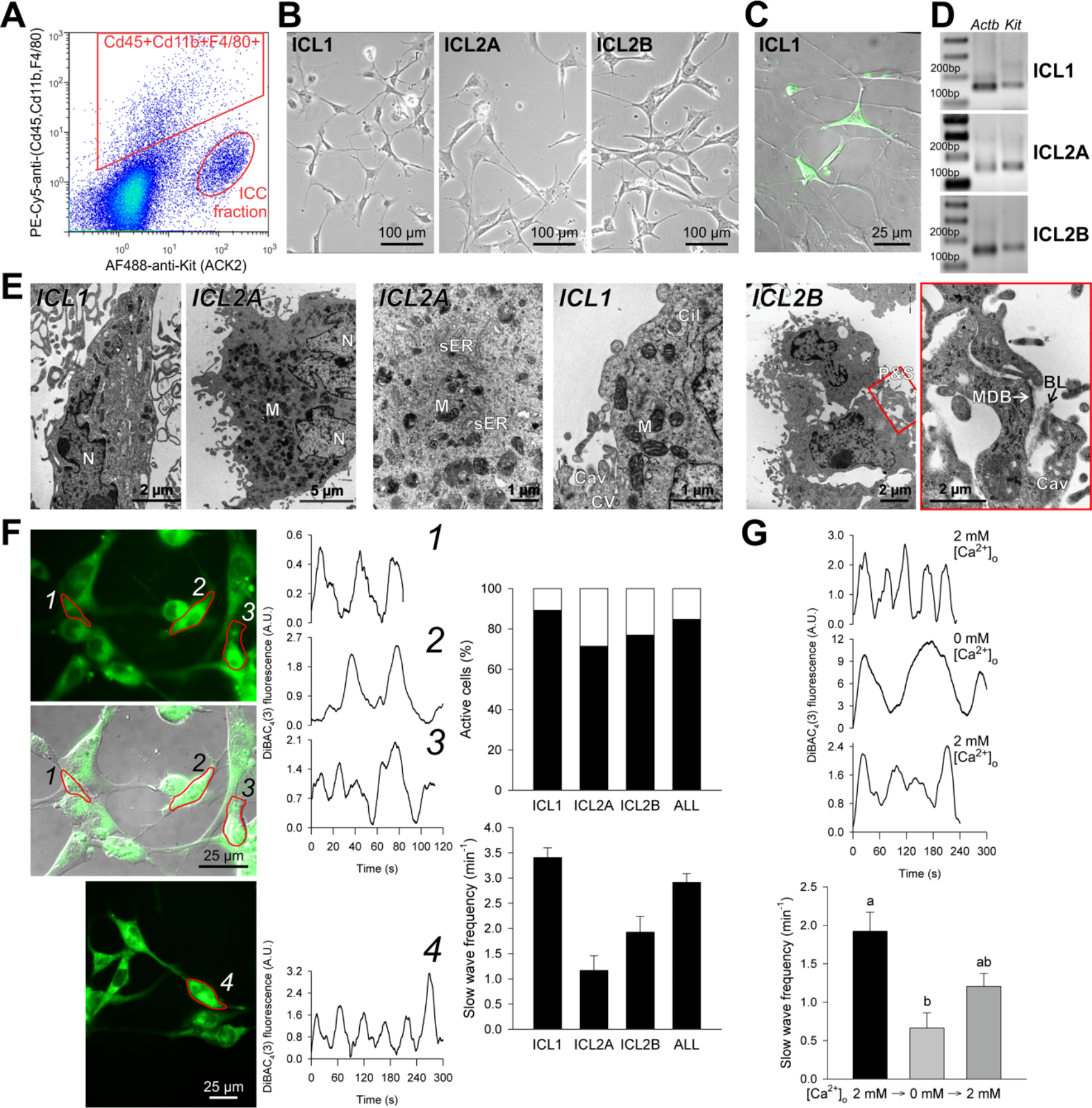
Conditionally immortalized ICC retain key ICC traits in early passages (<8). (**A**) Isolation of Kit^+^Cd45^−^Cd11b^−^F4/80^−^ cells from the gastric *tunica muscularis* of P12 *H-2K*^*b*^-*tsA58* mice by FACS following immunomagnetic depletion of Cd11b^+^Cd11c(Itgax)^+^ cells. PE-Cy5, phycoerythrin-cyanine 5. (**B**) Light microscopic morphology. (**C**) Kit-expressing cells detected by immunofluorescence (green) in the ICC-enriched ICL1 line. (**D**) *Kit* expression detected by reverse transcription-polymerase chain reaction (RT-PCR). (**E**) Ultrastructural features considered hallmarks of ICC: electron-dense nucleus (N) and cytoplasm, high density of mitochondria (M), abundant free ribosomes and intermediate filaments and smooth endoplasmic reticulum (sER) and numerous caveolae (Cav). Membrane-associated dense bands (MDB) with patchy basal lamina (BL), coated vesicles (CV) and cytoplasmic dense bodies were less frequent. The cells were interdigitating by peg-and-socket junctions (P&S) forming close appositions and small adherens junctions. Cil, primary cilium (9×2+0). The last image is an enlargement of the area outlined in red in the previous image. (**F**) Electrical oscillations reported by DiBAC_4_(3) fluorescence. Representative traces are from ICL2B cells identified by numbers in the fluorescent images. Bar graphs show the percentage of cells with phasic activity (top) and average (±s.e.m.) oscillatory frequency (bottom) in ICL1 (n=58), ICL2A (n=5), ICL2B (n=20) and all cells (n=83). (**G**) Reversible inhibition of phasic electrical activity by superfusion with Ca^2+^ - free medium. Representative traces are from ICL2B cells. Bar graph shows data (means±s.e.m.) combined from ICL1 (n=14), ICL2A (n=4) and ICL2B cells (n=5) recorded in 5 experiments before and 30 min after applying Ca^2+^-free medium; 1-h recovery data are from ICL2B cells. Groups not sharing the same superscript are different by Dunn’s multiple comparison (*P*<0.05) following Kruskal-Wallis ANOVA on ranks (*P*<0.001, *H*(2)=21.424). Panels **B-G** show results from cells maintained under conditions nonpermissive for tsTAg. A.U., arbitrary units. See also **Figure S2, Table S1**.

We then investigated the capability of the immortalized ICC to generate electrical pacemaker activity by monitoring the fluorescence of the voltage-sensitive dye bis-(1,3-dibutylbarbituric acid)trimethine oxonol (DiBAC_4_(3)) (Ordog et al., 2002). 83 of the 98 early-passage cells recorded displayed rhythmic activity at an average frequency of 2.92±0.35 min^−1^ (**Figure 2F**), which is typical of mouse antral ICC (Ordog et al., 2002). Consistent with the dependence of slow wave generation on extracellular calcium (Sanders et al., 2014), exposing ICL1, ICL2A or ICL2B cells to nominally calcium-free buffer for ~30 min reversibly reduced slow wave frequency (**Figure 2G**). Furthermore, similarly to cultured primary ICC (Ward et al., 2000), reducing the mitochondrial electrochemical gradient inhibited slow wave generation (**Figure S2C,D**). These observations indicate that conditionally immortalized ICC retain key morphological, ultrastructural, transcriptional and functional characteristics of ICC in early passages.

### Loss of ICC phenotype is recapitulated *in vitro*

Continuously maintained immortalized ICC gradually underwent phenotypic changes recapitulating the transition observed *in vivo*. Flow cytometry, immunofluorescent microscopy and Western blotting revealed a gradual reduction in extracellular then intracellular Kit protein levels, which first became evident by passage 7 (**Figure 3A-C, Figure S3A,B, Supplemental Experimental Procedures**). Growing the cells under conditions permissive, semipermissive or nonpermissive for the tsTAg or co-culturing with Kitl-expressing fibroblasts, which facilitated Kit expression in primary ICC (**Figure 3B**), did not affect Kit protein and mRNA expression (**Figure S3A,C,D** and not shown). After passage 15, Kit and Ano1 protein was barely or not detectable whereas Cd44 expression remained strong (**Figure S3A,B, Figure S3A**). The ICL lines maintained a dendritic morphology and Cd34 was not expressed regardless of culture duration.

**Figure 3.**
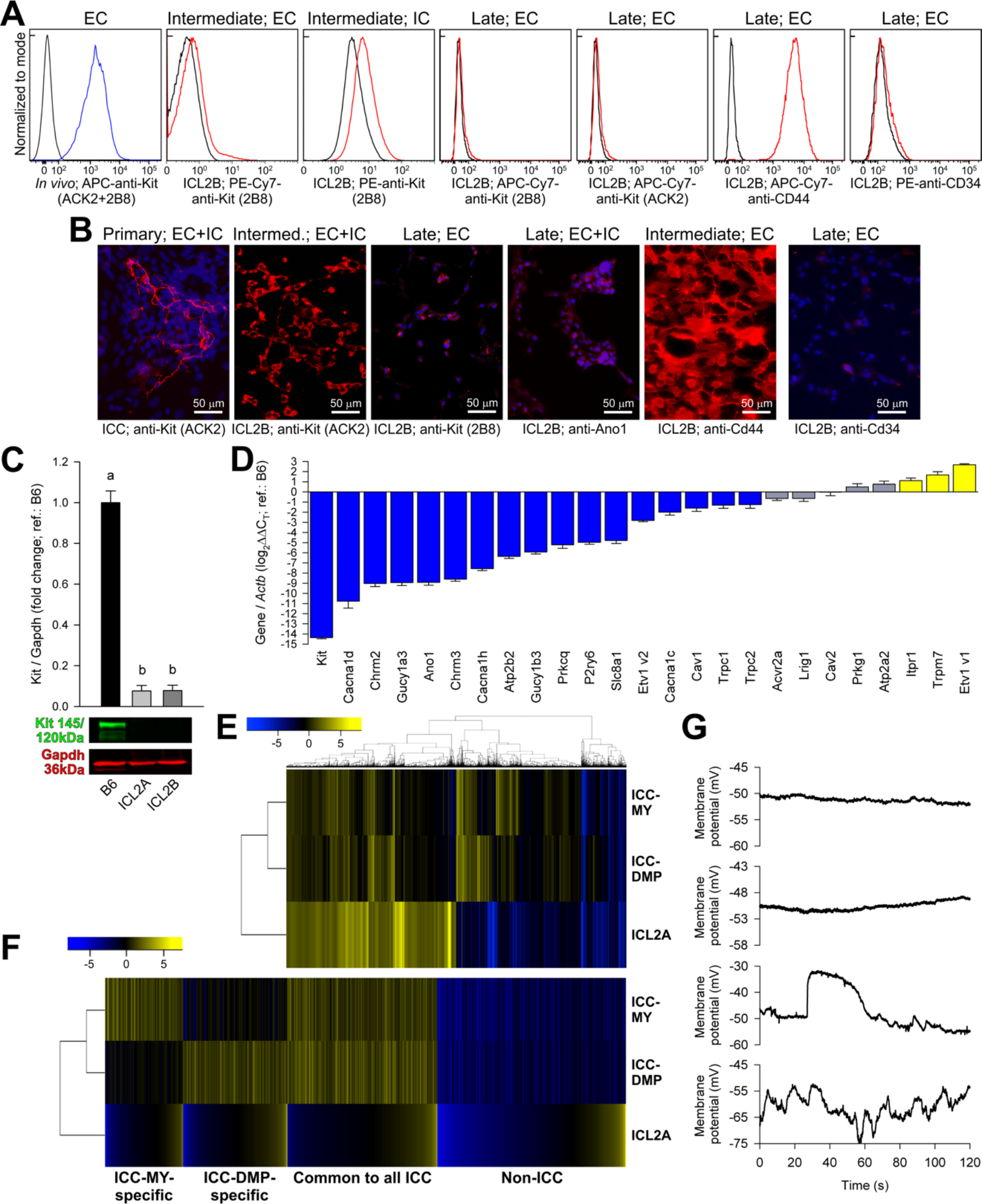
Conditionally immortalized ICC lose ICC phenotype with extended culturing. (**A,B**) Expression of indicated markers detected by (**A**) flow cytometry (permissive conditions) and (**B**) immunofluorescent microscopy (intensity-matched; red) in dissociated gastric muscles (*In vivo*), primary ICC co-cultured with Kitl-expressing fibroblasts and in intermediate-passage (7-14) and late-passage (27-31) ICL2B cells (nonpermissive). EC, extracellular immunolabeling; IC, intracellular immunolabeling. Blue pseudocolor: DAPI. Representatives of minimum 3 experiments are shown. (**C**) Reduced Kit expression detected by Western blotting in passage 20 ICL2A and passage 24 ICL2B cells (semipermissive) vs. C57BL/6J (B6) corpus+antrum muscles (means±s.e.m.; n=4/group). Groups not sharing the same superscript are different by Student-Newman-Keuls test (*P*<0.05) following Kruskal-Wallis nonparametric ANOVA (*P*=0.013, *H*(2)=7.449). (**D**) Downregulation of key ICC genes in ICL2A cells (semipermissive) by qRT-PCR. Comparative threshold cycle (log_2_ΔΔC_T_) values vs. B6 gastric muscles are shown (mean±s.e.m., n=3/gene). Blue: log_2_ΔΔC_T_≤−1; gray: −1<log_2_ΔΔC_T_<1; yellow: log_2_ΔΔC_T_≥1. (**E**) Heat map and hierarchical cluster analysis of 4,000 genes with the highest and lowest differential expression in ICL2A cells (semipermissive) vs. B6 corpus+antrum muscles detected by Affymetrix microarrays (n=3/group). Significant differential expression was defined as log2 fold change≤−1 or ≥1 with Benjamini-Hochberg *Q*<0.05. Differential gene expression in ICC-MY and ICC-DMP vs. source tissue was derived from GSE7809 (Chen et al., 2007). (**F**) Heat maps comparing differential expression of indicated ICC gene subsets between ICC-MY, ICC-DMP and ICL2A cells. (**G**) Patch clamp (current clamp) traces demonstrating lack of phasic activity (top two) and sporadic and irregular (bottom) slow wave activity. Representatives of 71 recordings from 20 late-passage cultures of 2 cell lines (ICL2A and ICL2B) are shown. See also **Figure S3, Supplemental Experimental Procedures** and **Supplemental Datasets 1 and 2**.

To further characterize the phenotype of ICL cells that lost Kit expression, we profiled mRNA expression by quantitative real-time RT-PCR (qRT-PCR) and Affymetrix Mouse Genome 430.2 microarray analysis in P23-26 ICL2A cells. By qRT-PCR performed using gastric *tunica muscularis* as reference, we detected consistent downregulation of 16 of 24 genes previously assigned to ICC based on expression profiling or genetic evidence (Chen et al., 2007; Chi et al., 2010; Cho and Daniel, 2006; Kondo et al., 2015; Peri et al., 2013; Sanders et al., 2014) including genes implicated in electrical slow wave generation and propagation (*Ano1, Cacna1h*) and neuromuscular neurotransmission of nitrergic (*Gucy1a3, Gucy1b3*) and cholinergic (*Chrm2, Chrm3*) signals (**Figure 3D**). By microarray analysis, we compared genes differentially expressed in ICL2A cells relative to gastric muscles to data previously generated in FACS-purified myenteric ICC (ICC-MY, which mainly function as electrical pacemaker cells) and deep muscular plexus-associated ICC (ICC-DMP, which are primary mediators of neuromuscular neurotransmission) using their source tissue as reference. Small intestinal ICC were used for this purpose because ICC functions are incompletely compartmentalized in the stomach (Hirst and Edwards, 2006). Initial heat map analysis and hierarchical clustering revealed profound differences between ICL2A cells and the two ICC classes (**Figure 3E**). Next, we generated gene sets specific for ICC-MY and ICC-DMP, and differential expression of these gene sets in ICL2A cells vs. gastric muscles was then determined (**Supplemental Dataset 1, Figure S3E**) and analyzed using heat maps (**Figure 3F**) and Ingenuity Pathway Analysis (IPA) (**Supplemental Dataset 2**). IPA of the 415 ICC-MY-specific genes indicated high representation of oxidative metabolism-related and calcium transport pathways, which have been implicated in ICC electrical pacemaking (Ward et al., 2000); and 141 (34%) of these genes including genes associated with oxidative phosphorylation were significantly downregulated in the ICL2A cells. The 552 ICC-DMP-specific genes were mainly associated with signaling pathways and 152 (28%) of these genes had significantly reduced expression in ICL2A cells including genes related to nitrergic, adrenergic and other neurotransmitter signaling pathways. The significant canonical pathways defined by the 555 genes common to all ICC were dominated by the immediate-early genes FBJ osteosarcoma oncogene (Fos) and Jun proto-oncogene and suppressor of cytokine signaling 3 (Socs3). In ICL2A cells, 137 (25%) of these common ICC genes were downregulated, and the related significant canonical pathways were also dominated by Fos and Jun.

To investigate whether the gene expression changes also impacted pacemaker function, we characterized the electrical activity of Kit^low/−^ ICL2A and ICL2B cells by performing patch clamp recordings in the current-clamp mode. Of the 71 cells recorded in 20 late-passage cultures, 42 (59%) did not display any spontaneous phasic activity and 22 cells (31%) only had sporadic (<1 min^−1^) slow waves (**Figure 3G, Figure S3F**). Seven cells (10%) showed phasic activity with a maximum frequency of 3.61 min^−1^. However, these events were highly arrhythmic relative to normal gastric slow waves (standard deviation of inter-event interval >30% of the mean; 95^th^ percentile of normal antral slow waves from ref.(Izbeki et al., 2010): 24%, n=67) (**Figure 3G**).

These findings indicate that similarly to post-ICC *in vivo*, continuously maintained conditionally immortalized ICC downregulate Kit expression without losing their dendritic phenotype. Furthermore, the *in-vitro* experiments also revealed loss of ICC-specific gene expression including genes related to electrical pacemaking and neurotransmitter signaling and a profound deterioration of electrical slow wave activity.

### ICC acquire the phenotype and function of Pdgfra^+^ interstitial cells *in vitro*

Because the majority of lineage-traced, Kit^low/−^ post-ICC expressed the key FLC marker Pdgfra *in vivo*, we next examined whether the clonally-derived ICL2A and ICL2B cells also gained FLC traits during their phenotypic transition into Kit^low/−^ cells. In late-passage cells, Pgdfra expression was readily detectable on the cell surface by flow cytometry and immunofluorescence, and Western blotting with a different antibody indicated robust overexpression relative to unfractionated gastric muscles (**Figure 4A-C, Figure S4A**). Strong *Pdgfra* mRNA expression was also detectable with published, validated primers (Peri et al., 2013) (**Figure 4D**) and with a second primer set validated in FACS-purified FLC using sorted ICC and ICC-SC as negative controls (**Figure S4E,F**).

**Figure 4.**
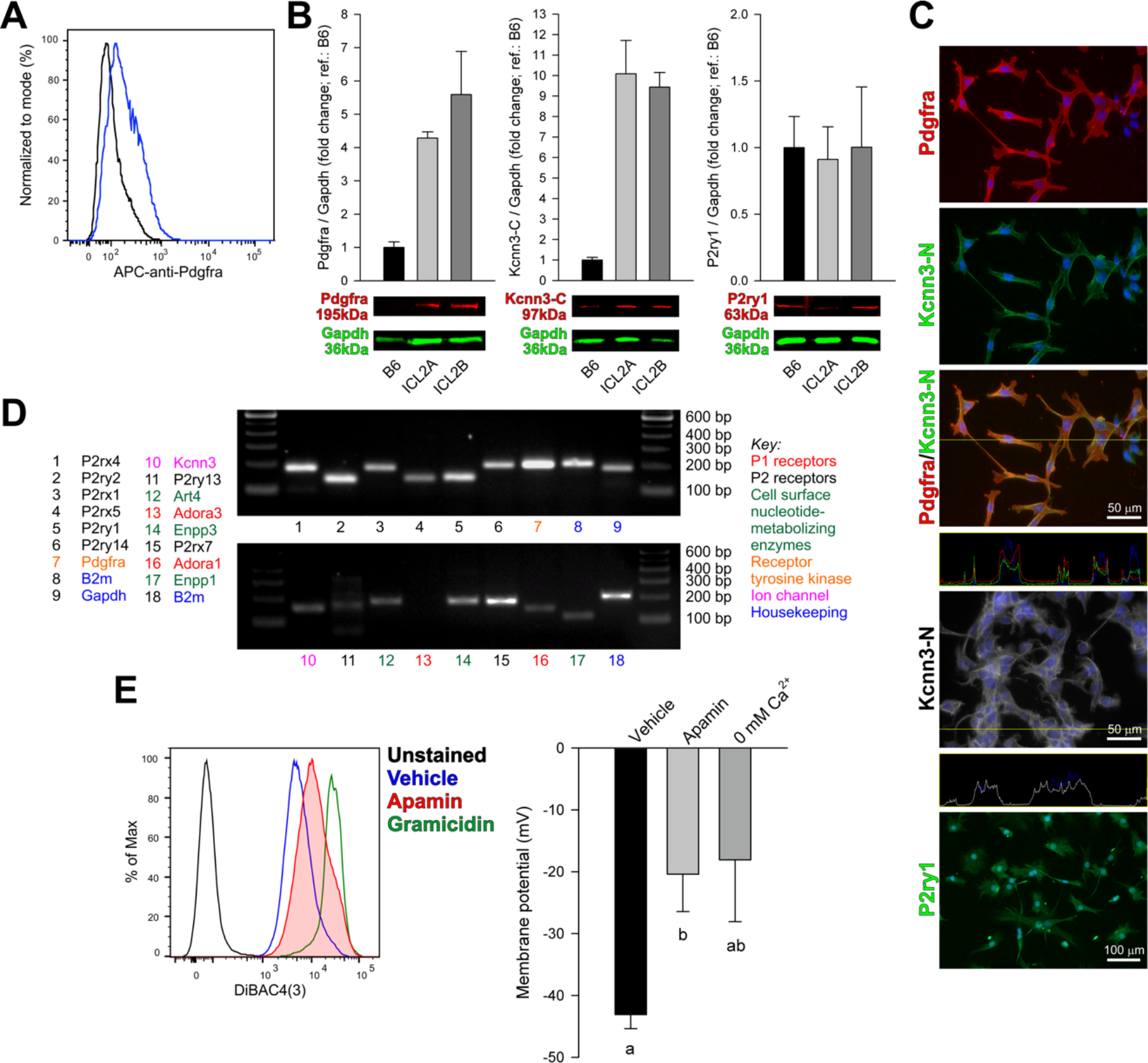
Conditionally immortalized Kit^low/−^ ICC gain FLC traits. (**A**) Pdgfra expression detected by flow cytometry in ICL2B cells (passage 31; semipermissive). A representative of 3 experiments is shown. (**B**) Expression (means±s.e.m.) of Pdgfra (*P*=0.05, *H*(2)=5.600, Kruskal-Wallis ANOVA on ranks, n=3/group), Kcnn3 (detected with an antibody against the C-terminus, Kcnn3-C; *P*=0.071, *H*(2)=5.422, n=3/group) and P2ry1 (*P*=0.99, *H*(2)=0.0201, n=5/group) analyzed by Western blotting in passage 18 ICL2A and passage 20 ICL2B cells (nonpermissive) vs. C57BL/6J (B6) corpus+antrum muscles. See uncropped blots in **Figure S4A-C**. (**C**) Detection of key FLC proteins by immunofluorecent microscopy. Top three panels: co-localization (orange) of Pdgfra (red) and Kcnn3 (detected with an antibody against the N-terminus, Kcnn3-N; green) in ICL2A cells (permissive). The fourth panel shows fluorescence profile along the yellow line in the merged image. Grayscale image: Kcnn3-N immunoreactivity in a different ICL2A culture. Fluorescence profile along the yellow line is shown in the panel below. Note fluorescence peaks associated with the cell membrane. The bottom panel demonstrates P2ry1 immunoreactivity in ICL2A cells (nonpermissive). Immunolabeling was most conspicuous in fine processes. Blue: DAPI. Representatives of 3 experiments are shown. (**D**) Detection of FLC-specific genes by RT-PCR using primers from ref.(Peri et al., 2013). Representative data from 3 experiments are shown. (**E**) Representative flow cytometry histograms and membrane potential values in ICL2A cells (permissive) in the presence or absence of 300 nM apamin or 0 mM extracellular Ca^2+^ (means±s.e.m.). Groups not sharing the same superscript are different by Dunn’s test (*P*<0.05) following Kruskal-Wallis nonparametric ANOVA, *P*=0.018, *H*(2)=8.000, n=6,6,3). See also **Figure S4, Supplemental Experimental Procedures, Supplemental Datasets 1 and 2**.

The FLC-specific ion channel Kcnn3 was colocalized with Pdgfra in the cell membrane by immunofluorescence employing a widely used antibody specific for the Kcnn3 N-terminus (Kcnn3-N) (**Figure 4C**), which we also found to exclusively label Pdgfra^+^ cells in murine gastric muscles (**Figure S4D**). Western blotting with a second antibody specific for the Kcnn3 C-terminus (Kcnn3-C) indicated increased expression in both ICL2A and ICL2B cells relative to gastric muscles (**Figure 4B**). Although *Kcnn3* mRNA levels detected with the published primers were relatively low (**Figure 4D**), expression could be confirmed using primers validated in FACS-purified FLC (**Figure S4F**), and small interfering RNA (siRNA)-mediated knockdown profoundly reduced Kcnn3-C protein detected by Western blotting (**Figure S4B**). The Kcnn3 channels were functional as both 0 mM extracellular Ca^2+^ and apamin, a highly selective inhibitor of Kcnn1, Kcnn2 and Kcnn3 channels (Kurahashi et al., 2011) depolarized ICL2A cells from a resting membrane potential of −43±2 mV (mean±s.e.m.) to −18±10 and −20±6 mV, respectively, as detected by DiBAC_4_(3) and flow cytometry (**Figure 4E**).

FLC express a variety of P1 and P2 purine/pyrimidine receptors and cell-surface nucleotide-metabolizing enzymes (Peri et al., 2013). Expression of P2ry1, the receptor most critical for enteric purinergic neuromuscular neurotransmission could be readily detected in ICL2A and ICL2B cells by immunofluorescence, Western blotting and RT-PCR using two different primer sets (**Figure 4B-D, Figure S4C,E**). Of the other 12 P1 and P2 receptors and extracellular nucleotide-metabolizing enzymes highly expressed in FLC, 11 were detected in ICL2A cells by RT-PCR (**Figure 4D**). Microarray analysis indicated that of the 1,000 genes downregulated in both ICC-MY and ICC-DMP (non-ICC genes), 152 (15.2%) were significantly upregulated in ICL2A cells (**Figure S4G**, **Supplemental Dataset 1**). These genes included enzymes of adenine and adenosine salvage pathways (**Supplemental Dataset 2**), further supporting the significance of purinergic signaling in the Kit^low/−^ ICL2A cells.

In contrast, similarly to their early-passage counterparts, Kit^low/−^ ICL2A and ICL2B cells did not express key smooth muscle markers (Myh11 and Tagln) by Western blotting (**Figure S4I, J**).

These results indicate that clonally selected, immortalized ICC fully recapitulate the fate of ICC that downregulate Kit expression during natural aging and acquire the phenotype and function of FLC.

### Genome-wide analysis reveals H3K27me3-mediated repression of ICC genes and H3K4me3-mediated activation of non-ICC genes *in vitro*

Because Kit is an established target of PRC2-mediated epigenetic repression in embryonic fibroblasts (Bracken et al., 2006), we examined whether PRC2-mediated H3K27me3 could underlie downregulation of Kit and other ICC genes in ICC transitioned into FLC *in vitro*. By microarray analysis, ICL2A cells expressed more mRNA for PRC2 core subunits Ezh2, Eed and Suz12 (Cao et al., 2002)) than ICC (**Figure 5A**, **Figure S5A**). Ezh2, Suz12 and H3K27me3 protein expression levels were higher in ICL2A and ICL2B cells than in unfractionated gastric muscles (**Figure 5B, Figure S5B-D**). In contrast, the PRC2-targeting gene *Jarid2* (Kaneko et al., 2014) was profoundly repressed. Of the 1,522 genes identified by microarray analysis as important for ICC-MY or ICC-DMP function, 182 (12%) were established PcG target genes (Bracken et al., 2006), and 78 (43%) of them were significantly repressed in the ICL2A cells relative to unfractionated gastric muscles (**Supplemental Dataset 1**). Chromatin immunoprecipitation (ChIP) experiments (**Figure 5C**; passage 22) indicated that the *Kit* transcription start site (TSS) was enriched with Ezh2 and H3K27me3 to a degree comparable to the TSS of the developmentally repressed PcG target *T* (brachyury), and neither contained the activating mark H3K4me3. The TSS of both *Kit* and *T* was also occupied by H3K9me3, a repressive mark established by Suv39h1, a member of a chromatin silencing complex distinct from PRC2. In contrast, the TSS of the constitutively active gene *Actb* was occupied by H3K4me3 but not Ezh2, H3K27me3 or H3K9me3.

**Figure 5.**
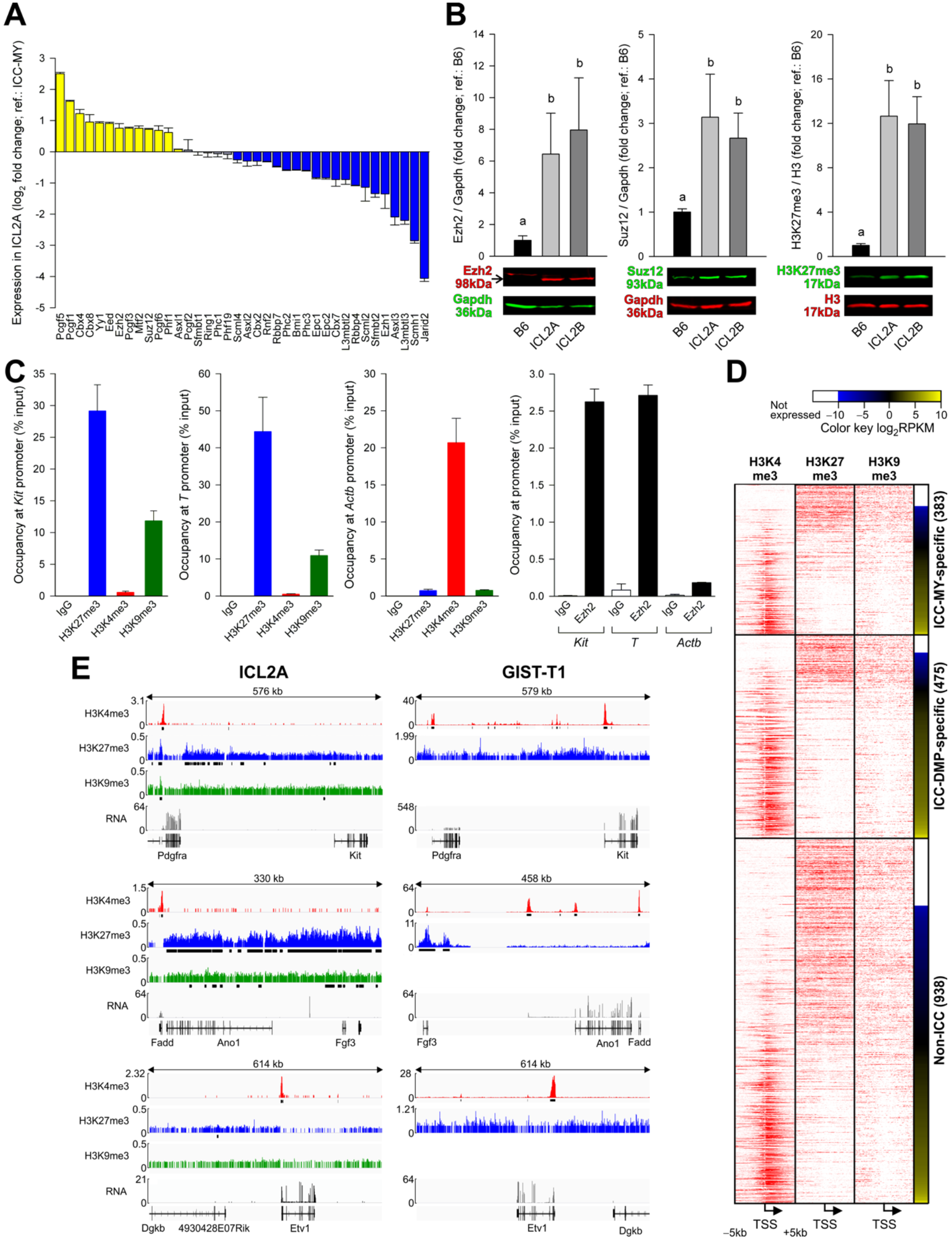
H3K27me3 occupies ICC genes after phenotypic transition into FLC. (**A**) Expression (means±s.e.m.) of PcG genes(Sauvageau and Sauvageau, 2010; Woo et al., 2010) in ICL2A cells relative to ICC-MY cells detected by microarray analysis (n=3/group; semipermissive). Yellow and blue indicate genes up- and downregulated in all ICL2A samples; gray: inconsistent change among replicates. (**B**) Expression (means±s.e.m.) of Ezh2 (*P*=0.009, *H*(2)= 9.380, Kruskal-Wallis ANOVA on ranks, n=5/group) and Suz12 (*P*=0.009, *H*(2)= 9.380, n=5/group) and H3K27me3 levels (*P*=0.001, *H*(2)= 13.365, n=7/group) analyzed by Western blotting in passage 20 ICL2A and passage 24 ICL2B cells (semipermissive) vs. C57BL/6J (B6) corpus+antrum muscles. Groups not sharing the same superscript are different by Student-Newman-Keuls test (*P*<0.05). See uncropped blots in **Figure S5B-D**. (**C**) ChIP on H3K27me3, H3K4me3, H3K9me3 and Ezh2 (n=2/target) in ICL2A cells (nonpermissive). Occupancy at the *Kit, T* (brachyury) and *Actb* TSS was determined by qPCR of ChIP DNA. (**D**) ChIP-sequencing on H3K4me3, H3K27me3 and H3K9me3 in ICL2A cells (nonpermissive). Tag density plots over TSS±5 kb displayed as heat maps for gene sets specific for ICC-MY, ICC-DMP and non-ICC and ranked by mRNA-sequencing. Representative data from 2 experiments are shown. RPKM, reads per kilobase exon per million mapped reads. Membership in gene subsets was slightly different from that reported for microarray data due to annotation differences (Affymetrix na33 for microarray and mm10 for RNA-seq data). (**E**) Genome browser tracks from the ChIP-sequencing (values: reads per million, RPM) and mRNA-sequencing (values: RPKM) experiments. GIST-T1: representatives of 2 independent ChIP-sequencing (on H3K4me3 and H3K27me3) and mRNA-sequencing experiments are shown. See also **Figures S5** and **S6, Supplemental Experimental Procedures** and **Table S2**.

Next, we performed ChIP- and mRNA-deep sequencing in passage 19-22 ICL2A cells to assess the genome-wide distribution of H3K27me3, H3K9me3 and H3K4me3 in relation to gene expression. Tag density plots for gene sets specific for ICC-MY, ICC-DMP and non-ICC and ranked by expression determined by mRNA-deep sequencing revealed strong inverse correlation between transcription and H3K27me3 occupancy and positive correlation between transcription and H3K4me3 occupancy in all gene sets (**Figure 5D, Figure 6A**). H3K9me3 occupancy was less correlated with gene transcription. Examination of the epigenomic profiles of 24 previously established ICC genes indicated that 12 genes lacking significant mRNA expression including *Kit* and *Ano1* did not possess H3K4me3 peaks at the TSS and were occupied by H3K27me3 and, to a much lesser extent, H3K9me3 peaks (**Figure 5E**, **Figure S6B-D**). In contrast, *KIT* and *ANO1* promoters had H3K4me3 peaks in KIT^+^ human GIST cells, the malignant counterpart of ICC (Hayashi et al., 2015). 10 genes including *Etv1* remained transcriptionally active and only had H3K4me3 peaks. In contrast, the TSS’s of transcriptionally activated non-ICC genes including *Pdgfra* were decorated with the activating H3K4me3 mark (**Figure 5D,E**, **Figure S6A**). Together, these data strongly support the hypothesis that in ICC transitioned into FLC *in vitro*, a significant subset of genes important for ICC function are transcriptionally silenced by Ezh2-mediated H3K27me3 and also indicate a role for H3K4me3 in the transcriptional activation of a subset of non-ICC genes.

**Figure 6.**
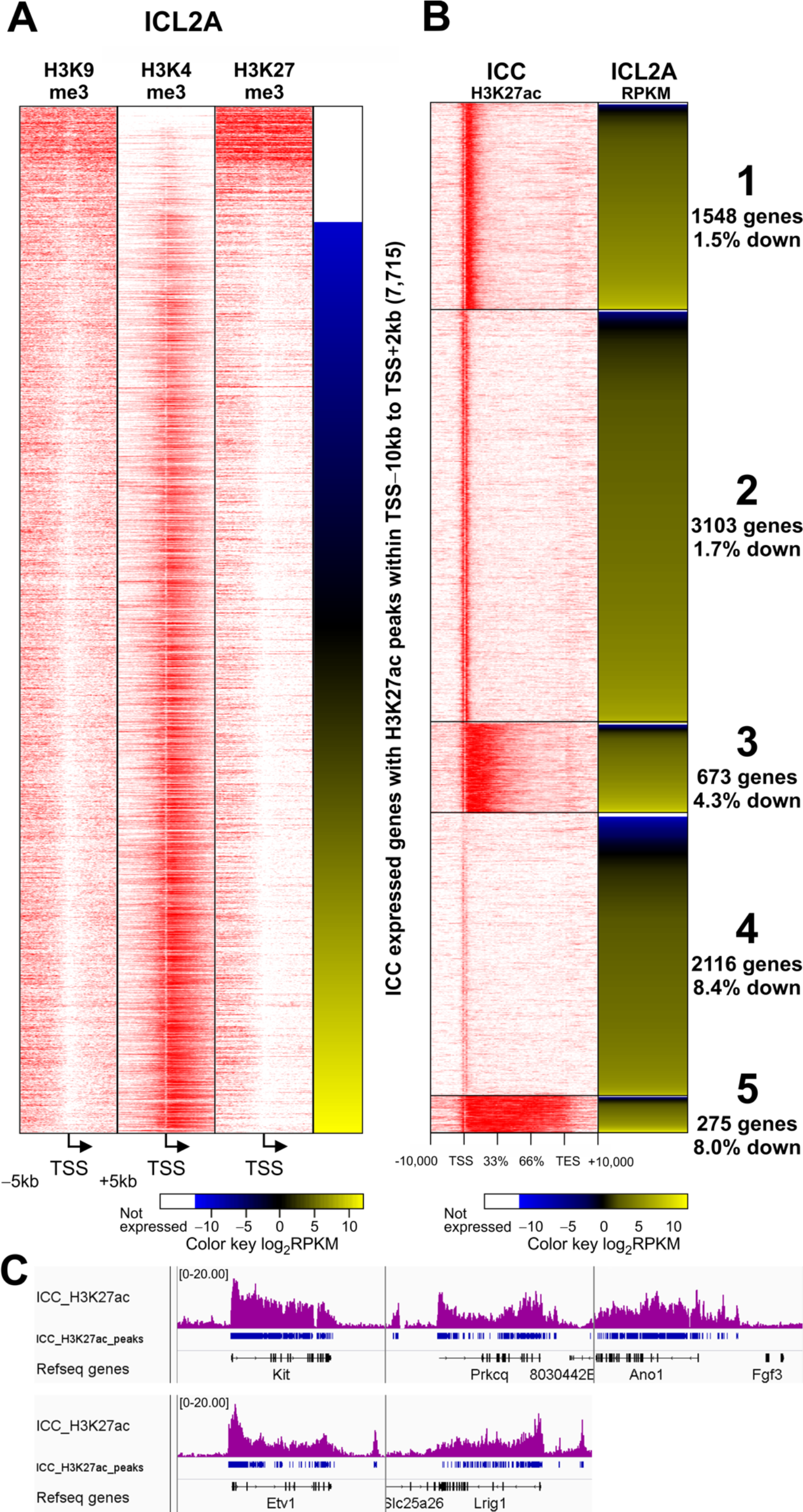
Pattern of H3K27 acetylation in ICC predicts ICC gene silencing in ICL2A cells. (**A**) H3K4me3, H3K27me3 and H3K9me3 ChIP-sequencing results from ICL2A cells (nonpermissive). Tag density heat maps (TSS±5 kb) for genes bearing significant H3K27ac peaks within TSS-10 kb and TSS+2 kb in FACS-purified small intestinal ICC are shown. Genes are ranked by mRNA expression in ICL2A cells (nonpermissive) determined by sequencing. Representative data from 2 experiments are shown. Note that the relationship between histone marks and gene expression is very similar to that seen in ICC-specific gene sets defined by microarray analysis (**Figure 5D**). (**B**) Patterns of TSS-10 kb to TES+10 kb occupancy by H3K27ac in FACS-purified small intestinal ICC determined by K-means clustering. Reproducibility of the ChIP-sequencing was verified by correlation analysis vs. colonic ICC (see **Table S2**). Five distinct clusters were identified. H3K27ac tag density heat maps are ranked cluster-wise according to gene expression in ICL2A cells determined by sequencing. Note that cluster 4 had the highest number and proportion of genes repressed in ICL2A. Cluster 5 had the second highest proportion but the overall number of these genes was small. % down values indicate the proportion of genes with RPKM<0.3 (Ramskold et al., 2009). (**C**) Genome browser tracks showing gene body-wide H3K27ac occupancy of *Kit, Prkcq, Ano1, Etv1* and *Lrig1* in ICC (values: RPM). These genes belonged to cluster 5. *Kit* and *Prkcq* met the criteria for transcriptional silencing (RPKM<0.3) and *Ano1* also had low expression.

### Genome-wide patterns of H3K27 acetylation predict transcriptional silencing during phenotypic switch of ICC into Pdgfra^+^ cells

The epigenomic landscape of ICC has not been characterized. Therefore, to understand why some ICC genes became preferentially silenced, we purified mG^+^ ICC from the intestines of tamoxifen-treated *Kit*^*CreERT2*/+^*;R26*^*mT-mG/mT-mG*^ mice by FACS and performed ChIP-sequencing on acetylated H3K27 (H3K27ac), a mark associated with active enhancers and promoters (Creyghton et al., 2010). 7,715 genes had significant H3K27ac peaks within TSS-10 kb to TES (transcription end site) +10 kb and were expressed in ICC by microarray analysis. This included 21 of the 24 previously established ICC genes (see **Figure 3D**). ICL2A mRNA-sequencing indicated that 306 of these genes were significantly downregulated including 5 ICC genes (*Kit, Cacna1d, Cacna1h, Atp2b2* and *Prkcq*). ChIP-sequencing showed increased occupancy of these genes by H3K27me3 and reduced binding by H3K4me3 (**Figure 6A**, **Figure S7A**, **Supplemental Dataset 1**). By K-means clustering of the 7,715 H3K27ac-occupied and expressed ICC genes, we discovered five distinct occupancy patterns (**Figure 6B**). Of these, cluster 4 representing genes with weak H3K27ac peaks contained the most downregulated genes (178; 8.4%; **Figure 6B**). Interestingly, the cluster containing proportionally the second most downregulated genes (22; 8.0%) was cluster 5 characterized by whole gene body occupancy. This group of genes included five key ICC genes (*Kit*, *Ano1*, *Etv1*, *Lrig1*, *Prkcq*; **Figure 6D**), which could be identified as super-enhancers by ROSE (Rank Ordering of SuperEnhancers) (Whyte et al., 2013) analysis. In ICL2A cells two of these genes (*Kit*, *Prkcq*) had mRNA expression meeting our predefined criteria for downregulation and *Ano1* was also repressed. These findings indicate that ICC genes with the weakest (cluster 4) and broadest (cluster 5) H3K27ac peaks are the most prone to transcriptional silencing.

### Pharmacological and genetic inhibition of Ezh2 reverses phenotypic loss of gastric ICC *in vitro* and *in vivo*

We next attempted to restore ICC gene expression and function in cells transitioned into FLC *in vitro* by siRNA-mediated and pharmacological inhibition of Ezh2. We employed adenosine dialdehyde (Adox) (Miranda et al., 2009), EPZ-6438 (Knutson et al., 2014) and GSK126 (McCabe et al., 2012) as small-molecule Ezh2 inhibitors. 2-week Adox treatment in ICL2A cells caused the re-expression of cell-surface Kit protein by flow cytometry, whereas 2-week treatment with the Ehmt2 (G9a) H3K9 methyltransferase inhibitor BIX-01294 or the histone deacetylase inhibitor suberanilohydroxamic acid (SAHA) and 4-week treatment with the DNA methyltransferase inhibitor RG108 had no effect (**Figure 7A**). Adox also increased Kit protein and mRNA while reducing H3K27me3 globally and over the *Kit, Ano1* and *Cacna1h* promoters in ICL2A cells (**Figure 7B, Figure S7B**). Kit protein was also increased by Adox in ICL2B cells (**Figure 7C**). EPZ-6438 also reduced global H3K27me3 and increased Kit protein expression (**Figure 7D**), and similar results were obtained with GSK126 (**Figure S7C**) and Ezh2 siRNA (**Figure 7E**). In ICL2A cells, 2-week Adox depolarized resting membrane potentials and increased the proportion of cells displaying electrical pacemaker activity from 10% (7/71) (see controls in **Figure 3G**) to 62.5% (10/16) (**Figure 7F**). Of these cells, 5 had regular activity not seen in vehicle-treated cells (standard deviation of inter-event interval <11.4% of the mean) occurring at a frequency of 3.99±0.09 min^−1^ (mean±s.e.m.) (**Figure 7F, Figure S7D,E**). The remaining cells displayed sporadic (<1 min^−1^; 2/16) or no activity (4/16). Thus, inhibition of Ezh2 can reactivate transcription of ICC genes including *Kit* by reducing TSS occupancy by H3K27me3 and restore Kit protein expression and rhythmic electrical activity.

**Figure 7.**
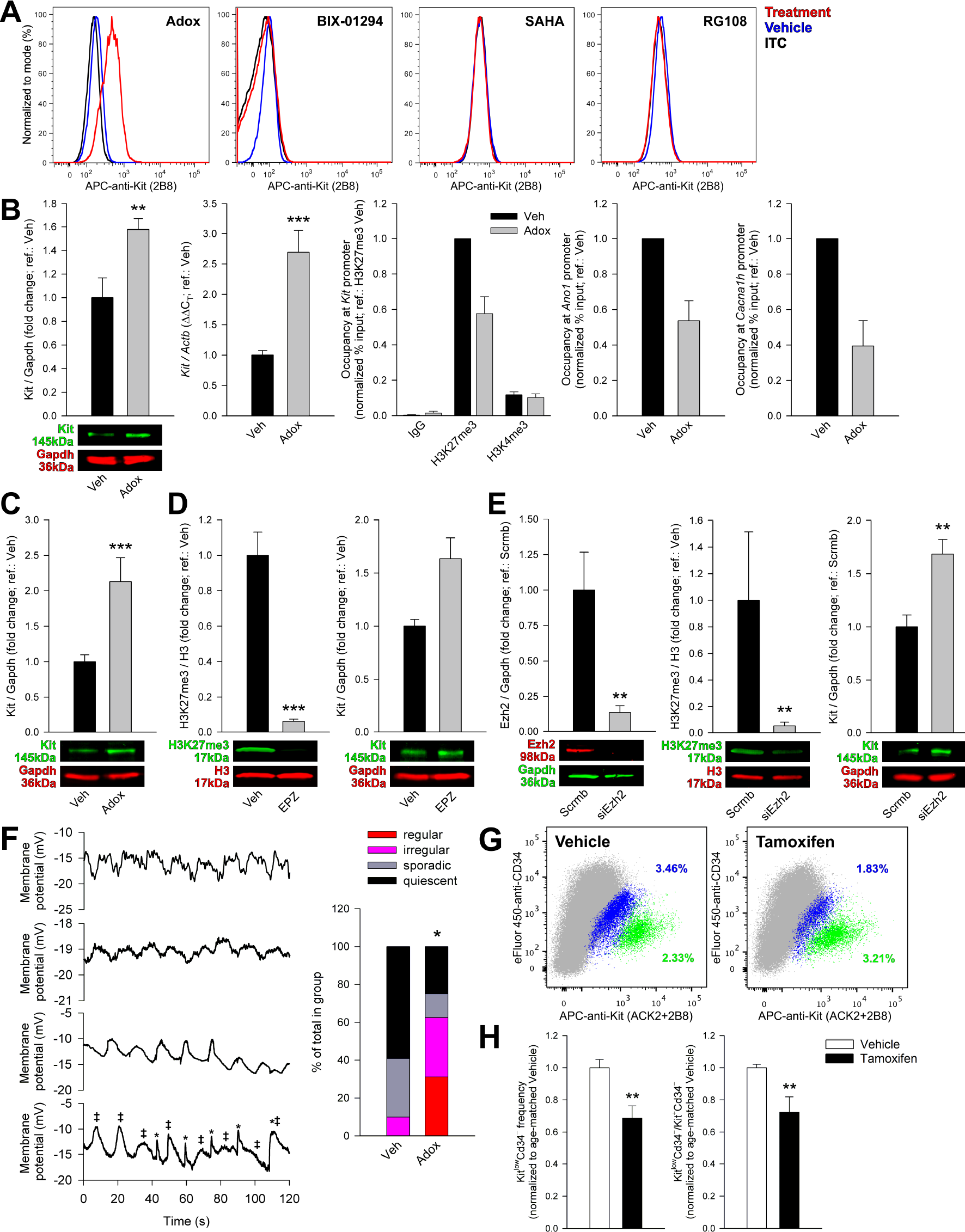
Ezh2 inhibition restores ICC phenotype and function. (**A**) Cell-surface Kit detected by flow cytometry in ICL2A cells treated with 2.5 μM adenosine dialdehyde (Adox) or 5 μM Ehmt2 inhibitor BIX-01294 or 1 μM histone deacetylase inhibitor suberanilohydroxamic acid (SAHA) for 2 weeks or 10 μM DNA methyltransferase blocker RG108 for 4 weeks. (**B**) In passage 19-39 ICL2A cells, 2.5 μM Adox increased Kit protein (Western blotting; 2 weeks; permissive; n=6/group) and *Kit* mRNA (qRT-PCR; permissive; 4 days; n=6/group) while reducing H3K27me3 occupancy of the *Kit, Ano1* and *Cacna1h* TSS without affecting H3K4me3 binding (ChIP; semipermissive; 4 days; n=3/group/target). Means±s.e.m. are shown. **, *P*=0.026, ***, *P*=0.002. Veh, vehicle; IgG, non-targeting antibody. (**D**) In passage 11 ICL2B cells (permissive), 2.5 μM Adox (2 weeks) increased Kit protein. ***, *P*=0.002, n=6/group. (**D**) In passage 18 ICL2A cells (permissive), 2-week treatment with 500 nM EPZ-6438 (EPZ) reduced H3K27me3 and increased Kit protein (n=9/group/experiment; ***, *P*<0.001). (**E**) In passage 11 ICL2A cells (semipermissive), 2-week exposure to siRNA targeting Ezh2 (siEzh2) reduced Ezh2 protein (**,*P*=0.004, n=6/group) and H3K27me3 levels (**,*P*=0.013, n=9/group) and increased Kit protein (**, *P*=0.009, n=6/group). Scrmb, scrambled siRNA. (**F**) Top: Representative patch clamp (current clamp) traces demonstrating rhythmic slow wave activity. Bottom trace: partially interfering events from two unentrained pacemakers (labeled with ‡ and *) likely representing electrically coupled cells. Representatives of 16 recordings from 4 cultures of ICL2A cells are shown. Bottom: proportional representation of activities in vehicle-treated (n=71) and Adox-treated (n=16) cells; see text for definitions. (**G-H**) Representative and quantitative flow cytometry data from the non-hematopoietic (HP^−^) fraction of the gastric corpus^+^antrum *tunica muscularis* of *Kit*^*CreERT2*/+^*;Ezh2*^*fl/fl*^ mice treated with tamoxifen (n=7) or vehicle (n=5) between P77-151. ICC and post-ICC were quantified 9 weeks after the first injection. (**G**) Representative dot plots (100,000 HP^−^ cells; gray) showing reduced HP^−^Kit^low^Cd34^−^ post-ICC (blue) and increased HP^−^Kit^+^Cd34^−^ ICC (green). (**H**) Reduced frequency of post-ICC (left; **, *P*=0.018) and post-ICC/ICC ratios (right; *P*=0.03) signifying reduced ICC-to-post-ICC transition. See also **Figure S7**.

With *in-vitro* data suggesting that Ezh2 mediates the ICC to post-ICC transition, we next aimed to reverse ICC loss by deleting the *Ezh2* gene in Kit-transcribing cells in adult *Kit*^*CreERT2*/+^*;Ezh2*^*fl/fl*^ mice. Seven *Kit*^*CreERT2*/+^*;Ezh2*^*fl/fl*^ mice received tamoxifen and 5 vehicle treatment between P77-151. HP^−^Kit^+^Cd34^−^ ICC and HP^−^Kit^low^Cd34^−^ post-ICC were quantified 9 weeks after the first injection by flow cytometry. None of the mice expressed any outward phenotype. Cre-mediated Ezh2 knockout resulted in significantly reduced post-ICC frequency and post-ICC/ICC ratios signifying reduced ICC-to-post-ICC transition (**Figure 7G,H**).

## DISCUSSION

*In-vivo* transdifferentiation is a well-characterized form of cellular plasticity in invertebrates and amphibians but only few *bona-fide* examples have been reported in mammals and even fewer under physiological conditions (Chera et al., 2014; Merrell and Stanger, 2016; Yanger et al., 2013). Furthermore, the underlying epigenetic mechanisms remain largely unexplored. Here, using “pulsed” genetic lineage tracing in pre-weaning mice we show that during the first 3-5 months of life, the natural fate of ~16% of Kit^+^ gastric ICC is transdifferentiation into Pdgfra^+^ FLC, a functionally distinct mesenchymal cell type (Iino et al., 2009; Kurahashi et al., 2008; Peri et al., 2013; Sanders et al., 2014). The RTK switch observed *in vivo* was faithfully recapitulated in clonal immortalized ICC lines maintained without specific stimulation of the ICC RTK Kit. While losing expression of genes important for established ICC functions and also electrical slow wave activity, these cells gained expression of 14 of 15 genes previously assigned to FLC including *Pdgfra, P2ry1* and several P1 and P2 purine/pyrimidine receptors and cell-surface nucleotide-metabolizing enzymes, as well as *Kcnn3* (Kurahashi et al., 2011; Peri et al., 2013), which conferred apamin-sensitive hyperpolarization of post-ICC membrane potentials. The ICC-to-FLC transdifferentiation occurred as a part of a major age-related remodeling of the gastric neuromuscular apparatus involving another 12% of ICC losing their phenotype and function without gaining FLC traits (post-ICC) and the death of yet another 21%. We found no evidence of ICC reverting to ICC-SC during this process as post-ICC, the cell type from which FLC probably differentiate, could be distinguished from ICC-SC (Bardsley et al., 2010; Lorincz et al., 2008). Importantly, we also demonstrate that transcriptional repression of key ICC genes including *Kit* occurred by Ezh2-mediated H3K27 trimethylation, while transcriptional activation of non-ICC genes involved increased H3K4 trimethylation. The ICC genes’ propensity for repression could be predicted by their patterns of H3K27ac occupancy in ICC. Our findings also indicate that repression of ICC transcriptional programs is likely the primary component of the ICC-to-FLC transdifferentation as ICC identity and pacemaker activity could be recovered by Ezh2 inhibition, knockdown or genomic deletion even after the phenotypic transition had taken place. Besides demonstrating an instance of phenotypic conversion between two specialized cell types during mouse postnatal development, our studies identify an epigenetic mechanism, specifically, Ezh2-mediated H3K27 trimethylation, playing a key role in physiological mammalian transdifferentiation.

ICC are prone to depletion in numerous GI disorders (Farrugia, 2008; Ordog, 2012; Sanders et al., 2014) and their numbers decline substantially with age (Gomez-Pinilla et al., 2011; Izbeki et al., 2010), setting up the neuromuscular apparatus for decompensation and failure (e.g., gastroparesis) in response to metabolic and other challenges such as infections, stress, or even a meal (Ordog, 2012). In humans, ICC decline at a rate of ~13% per decade of life between 25 and 92 years of age (Gomez-Pinilla et al., 2011). Our present data in mice indicate a more rapid time course in early life leading to a loss similar to that seen in progeric *klotho* mice (Izbeki et al., 2010) by P107-160 (57%). We now show that part of this age-related ICC depletion is transdifferentiation resulting in approximately 3% of the FLC being ICC-derived at the end of the study. While this phenotypic switch may in part explain the lack of susceptibility of FLC to injury e.g. in diabetic and idiopathic gastroparesis (Grover et al., 2012), this small gain in FLC and purinergic inhibitory neuroeffector functions is likely much less significant functionally than the ICC loss *per se*. ICC depletion can lead to GI motor dysfunction through deterioration and loss of electrical slow waves required for normal phasic contractile activity and orderly peristalsis (Huizinga et al., 2014; Klein et al., 2013; Sanders et al., 2014) and impaired excitatory cholinergic and inhibitory nitrergic neuromuscular neurotransmission (Groneberg et al., 2015; Klein et al., 2013; Lies et al., 2014; Sanders et al., 2014). The potential translational significance of the detected ICC transdifferentiation is in their reversibility: post-ICC and ICC-derived FLC appear to constitute a reserve from which functioning ICC can be restored using available pharmacological agents, raising hopes for novel therapeutic options for GI neuromuscular disorders which, despite their high prevalence, lack effective therapy (Ordog, 2012) and cost billions of dollars in health care spending each year (2009).

Comparing transcriptional profiles of ICC transitioned into FLC *in vitro* to selectively harvested ICC-MY and ICC-DMP identified downregulation of genes important for ICC identity (*Kit, Prkcq, Fos, Jun, Socs3*), pacemaker function (*Ano1*, *Cacna1h*, *Atp2b2*, *Slc8a1* and genes of oxidative metabolism), and neuromuscular neurotransmission mediated by nitric oxide (*Gucy1a3*, *Gucy1b3*) and acetylcholine (*Chrm2*, *Chrm3*) (Chen et al., 2007; Sanders et al., 2014). These changes were accompanied by loss or deterioration of electrical pacemaker activity. In contrast, caveolin mRNA remained detectable. Since FLC do not typically have caveolae, presence of this organelle may distinguish ICC-derived FLC from FLC developed from embryonic precursors (Tamada and Kiyama, 2015). Downregulation of soluble guanylyl cyclase isoforms *Gucy1a3* and *Gucy1b3* may also distinguish between these cells since these genes are repressed in post-ICC but expressed in FLC (Groneberg et al., 2015). ChIP-sequencing indicated that accumulation of H3K27me3, established by the PRC2 histone methyltransferase Ezh2, was mainly responsible for the repression of ICC genes. Pharmacological and siRNA-mediated blockade or Cre-mediated *in-vivo* genomic inactivation of Ezh2 increased Kit expression promoting ICC re-differentiation and the recovery of electrical pacemaker activity *in vitro*. However, the slow waves were small likely due to the depolarized diastolic potentials (Sanders et al., 2014). Stimulating Kit signaling by supplying membrane-bound Kitl to cells with recovered Kit expression may be necessary to completely restore the expression of genes required for normal slow wave activity. More robust Kit signaling may in turn facilitate functional recovery by increasing the levels of Etv1 transcription factor, which in GIST targets several genes related to ICC function including *ANO1, CACNA1H, CACNA1D, CHRM2, PRKG1* and *PRKCQ* (based on ChIP-sequencing data from GSE22852 (Chi et al., 2010)). While we found *Etv1* to be transcribed in ICL2A and ICL2B cells, Etv1 protein was undetectable (not shown) and may not have been adequately upregulated following Ezh2 inhibition in the absence of ligand-induced Kit activation.

We found that approximately 30% of ICC-specific genes became silenced during transition to FLC. To get insight into why some ICC genes became preferentially downregulated, we analyzed their H3K27ac occupancy in freshly purified ICC. The majority of downregulated genes had weak H3K27ac over or upstream the TSS. This is not surprising since genes with weak histone acetylation and low transcription are more prone to repression (Kaneko et al., 2013). However, 22 genes that became downregulated including the key ICC genes *Kit* and *Prkcq* had strong H3K27ac coverage of their entire gene body. In fact, genes with strong H3K27ac occupancy were almost as likely as genes with weak H3K27ac peaks to become silenced. Preliminary analysis of these 22 genes indicated that 5 (*Kit, Prkcq, Gpr20, Ifitm1*, and *Mab21l2*) are Etv1 targets (Chi et al., 2010) and contain super-enhancers in ICC. Thus, silencing of key ICC genes is likely to begin with enhancer decommissioning due to reduced Etv1 binding and histone acetylation. Loss of transcriptional activation then sets the stage for Ezh2-mediated repression, which takes over as the primary gatekeeper of re-activation. Consistent with this concept, we found that in ICC-derived FLC, histone deacetylase inhibition was already ineffective at restoring Kit expression.

It remains unclear why these mechanisms are triggered during aging or disease and why other genes including genes related to FLC function become aberrantly de-repressed. A strong candidate is insufficient activation of Kit by Kitl from the microenvironment (Hayashi et al., 2013; Izbeki et al., 2010; Ordog, 2012), which sets in motion the epigenetic cascade of enhancer loss and H3K27me3-mediated repression. Reduced Kit signaling can interfere with transcriptional activation by facilitating Etv1 degradation (Chi et al., 2010; Hayashi et al., 2015) and by reducing histone acetylation (Huang and Chen, 2005). Furthermore, loss of Etv1 may also influence PRC2 recruitment via the Jumonji family protein *Jarid2* (Kaneko et al., 2014), an Etv1 target (Chi et al., 2010) we found to be profoundly repressed in ICL2A cells. De-repression of FLC genes may occur under the influence of Pdgfra, which our present and previous data (Hayashi et al., 2015) indicate to be in inverse relationship with Kit in non-malignant cells. Pdgfra signaling alone does not appear to be sufficient to maintain Etv1 protein levels and thus rescue ICC genes from repression but could stimulate expression of the FLC gene set by mechanisms that are yet to be determined. Interestingly, we found 5 of the 15 FLC genes to bear H3K27ac peaks in ICC (*Art4, Enpp1, P2rx1, P2rx4* and *P2rx7*) suggesting that some of these genes may be poised for activation even in ICC. These results further support the notion that Kit^+^ ICC and Pdgfra^+^ FLC are closely related cell types.

## EXPERIMENTAL PROCEDURES

Further details can be found in the **Supplemental Experimental Procedures**.

### Animal models

Experiments were performed in accordance with the National Institutes of Health Guide for the Care and Use of Laboratory Animals. All protocols were approved by the Institutional Animal Care and Use Committee of the Mayo Clinic. *Kit*^*CreERT2*/+^ mice (background: C57BL/6) and Homozygous *Gt(ROSA)26Sor*^*im4(ACTB-tdTomato-EGFP*^ (*R26*^*mT-mG/mT-mG*^) reporter micewere kindly provided by D. Saur. *Ezh2*^*fl/fl*^ mice (background: C57BL/6) carrying *loxP* sites between *Ezh2* exons 15-16 and 19-20, flanking the essential SET domain encoded by exons 16-19 were obtained from R.A. Urrutia. *Kit*^*CreERT2*/+^*;R26*^*mT-mG/mT-mG*^ and *Kit*^*CreERT2*/+^*;Ezh2*^*fl/fl*^ mice were generated in our breeding program. Homozygous *H-2K^b^-tsA58* transgenic mice (Immortomice^®^ CBA;B10-Tg(H2K^b^-tsA58)6Kio/Crl) harboring the temperature-sensitive simian virus 40 tsA58 mutant large T antigen (tsTAg) were from Charles River Laboratories (Wilmington, MA). C57BL/6J, and NOD/ShiLtJ mice were purchased from The Jackson Laboratory (Bar Harbor, ME) and BALB/c mice from Harlan Laboratories (Madison, WI). Mice were housed maximum 5/cage in the Mayo Clinic Department of Comparative Medicine Guggenheim Vivarium under a 12 h light/12 h dark cycle. Both male and female mice were used in all experiments. None of the mice were used in any previous experiments.

### Induction of Cre-mediated recombination *in vivo*

Cre-mediated recombination was induced with tamoxifen (Sigma-Aldrich, St. Louis, MO) injected intraperitoneally once daily for 3 consecutive days at a dose of 0.075 mg/g body weight in peanut oil vehicle (Sigma Aldrich) containing 10% ethanol (3.75 μL/g body weight). For lineage tracing, 12 *Kit*^*CreERT2*/+^*;R26*^*mT-mG/mT-mG*^ mice underwent tamoxifen treatment between postnatal days (P) 8-10. Six *Kit*^+/+^*;R26*^*mT-mG/mT-mG*^ mice were treated with vehicle only and 4 *Kit*^*CreERT2*/+^*;R26*^*mT-mG/mT-mG*^ and strain-matched wild-type mice were left untreated. Experiments were performed at postnatal day (P) 11 (n=7) and between P107-P160. For validation of in-vivo lineage tracing, see **Supplemental Experimental Procedures**. For cell sorting, 14 *Kit*^*CreERT2*/+^*;R26*^*mT-mG/mT-mG*^ mice were treated with tamoxifen starting on P28-45 and killed 24 h after the last injection. Seven *Kit*^*CreERT2*/+^*;Ezh2*^*fl/fl*^ mice received tamoxifen treatment between P77-P151. Five *Kit*^*CreERT2*/+^*;Ezh2*^*fl/fl*^ mice were treated with vehicle on the same days. Experiments were performed 9 weeks after the first injection.

### Multi-parameter flow cytometry analysis of cells of the interstitial cell of Cajal (ICC) lineage

ICC and related cells (see **Figure S1A,B**) were identified in the hematopoietic marker-negative fraction of dissociated mouse gastric corpus+antrum *tunica muscularis* using previously published protocols with modifications. A detailed protocol and other experimental details are provided in the **Supplemental Experimental Procedures**.

### ICC purification by fluorescence-activated cell sorting (FACS)

In 14 adult, tamoxifen-treated *Kit*^*CreERT2*/+^*;R26*^*mT-mG/mT-mG*^ mice, ICC in the small intestines (jejunum+ileum) and the colon were identified as mG^+^HP^−^ cells and sorted by FACS. DNA-protein complexes in the dissociated cells were cross-linked using fresh 1% formaldehyde (Thermo Fisher Scientific, Waltham, MA) for 10 min at room temperature followed by glycine quenching (125 mM, 5 min, room temperature). Cells were stored at 4 °C overnight, then labeled for hematopoietic markers (Cd45, Cd11b and F4/80) and sorted using a BD Biosciences FACSAria II instrument configured to match the LSR II cytometer. Gastric ICC, ICC stem cells (ICC-SC) and fibroblastlike cells (FLC) were purified from BALB/c mice as Kit^+^Cd44^+^Cd34^−^, Kit^low^Cd44^+^Cd34^+^ and Kit^−^ Cd44^−^Cd34^−^Pdgfra^+^ cells, respectively.

### Conditionally immortalized ICC

ICL2A and ICL2B cells were isolated from gastric corpus+antrum *tunica muscularis* tissues of two homozygous, 12-day-old Immortomice by fluorescence-activated cell sorting (FACS) using a previously published and validated approach and antibodies (see **Supplemental Experimental Procedures**).

### Primary ICC cultures

Primary ICC were cultured in the presence of stromal cells derived from the fetal hematopoietic microenvironment of Kitl-deficient *Kit*^*Sl/Sl-4*^ mice and genetically modified to express full-length murine Kitl (Kitl^248^) as described (Rich et al., 2003).

### Immunocytochemistry and immunohistochemistry

Cultured cells were fixed with 4% paraformaldehyde, permeabilized with Triton X-100 (Sigma-Aldrich) and immunolabeled as described previously (Bardsley et al., 2010) (see antibodies and detailed protocol in **Supplemental Experimental Procedures**). Cells were examined using a Nikon (Melville, NY) Eclipse TS-100F fluorescent microscope equipped with Hoffman Modulation Contrast (Glen Cove, NY) objectives and a Jenoptik (Brighton, MI, USA) MFcool CCD digital camera. Postacquisition manipulation of fluorescent images was restricted to adjustments to brightness and contrast and assignment of pseudocolor, which were performed in each optical channel separately.

### Gene expression analysis by reverse transcription-polymerase chain reaction (RT-PCR)

Quantitative real-time RT-PCR (qRT-PCR) and end-point RT-PCR were performed using performed using previously published methods (Izbeki et al., 2010) and specific, intron-spanning primers (Thermo Fisher and Roche Applied Science, Indianapolis, IN) (**Supplemental Experimental Procedures** and ref.(Peri et al., 2013)). For qRT-PCR the cDNA was amplified on a Bio-Rad (Hercules, CA) CFX96 or a Roche LightCycler 480 (Roche Applied Science) realtime PCR detector using the SYBR GreenER qPCR SuperMix (Thermo Fisher). For end-point RT-PCR, cDNA was amplified using a Bio-Rad C1000 Touch Thermal Cycler and REDTaq ReadyMix PCR Reaction Mix (Sigma-Aldrich).

### Transmission electron microscopy

Cells were lifted by trypsinization and immersion-fixed for 7 days in a fixative containing 2 % paraformaldehyde and 2% glutaraldehyde (Sigma-Aldrich) in 0.1 M cacodylate buffer (pH 7.3). After rinsing for 60 min in 0.1 M cacodylate buffer (pH 7.3), cells were postfixed in 1% OsO_4_ in 0.1 M phosphate buffer for 1 h, dehydrated in alcohol, and embedded in Epon 812 R (Merck, Kenilworth, NJ). Ultrathin (50-70 nm) sections were cut, mounted on copper grids, contrasted with uranyl acetate and lead citrate and examined using a Philips Morgagni 268 electron microscope (Philips, Eindhoven, Netherlands) at 80 KV.

### Analysis of electrical pacemaker activity by fluorescent microscopy

Electrical activity of conditionally immortalized ICC was assessed by monitoring the fluorescence of bis-(1,3-dibutylbarbituric acid)trimethine oxonol (DiBAC_4_(3); 1 μmol/L; Thermo Fisher), a voltage-sensitive dye, as described (Ordog et al., 2002), with the following modifications. Imaging was performed using a Nikon Eclipse E600FN microscope equipped with a Fluor 60×, 1.00 NA WI DIC objective. Fluorescence excitation at 494±7.5 nm was generated with a Polychrome IV monochromator (FEI, TILL Photonics, Gräfelfing, Germany). Images were acquired using a TILL Photonics Imago QE camera and TILLvisION 4.1 software. Triflouromethoxyphenylhydrazone (FCCP) and antimycin A were from Sigma-Aldrich.

### Characterization of gene expression by flow cytometry

Expression of key markers in conditionally immortalized ICC was analyzed using established techniques (Bardsley et al., 2010). Cells labeled with isotype control antibodies were used as reference. Flow cytometry was performed on a BD Biosciences LSR II flow cytometer and analyzed using FlowJo 10 software (Tree Star). Antibodies and cytometer configuration are provided in the **Supplemental Experimental Procedures**.

For assessment of surface and intracellular Kit expression simultaneously, unfixed cells were stained with PE-Cy7-anti-Kit, clone 2B8 for 30 minutes at 4 °C and washed. Cells were resuspended in PBS/Nonidet P40 permeabilization buffer (BioSure, Grass Valley, CA) and incubated for 10 minutes at room temperature. Permeabilized cells were washed and labeled with anti-mouse Cd16/32 antibody (Fc block) for 10 minutes at 4°C, stained with the same antibody (2B8) labeled with PE for 30 minutes at 4°C. Flow cytometry was performed using a Beckman Coulter (Brea, CA) XL/MCL instrument.

### Western blotting

Tissue and cell lysates were prepared and subjected to sodium dodecyl sulfate-polyacrylamide gel electrophoresis and immunoblotting as described previously (Hayashi et al., 2013) (see antibodies in **Supplemental Experimental Procedures**). Specificity of the Kcnn3 band obtained with the antibody recognizing the C-terminus was verified by small interfering RNA (siRNA)-mediated knockdown. Post-acquisition manipulation of fluorescent images was restricted to adjustments to brightness and contrast and assignment of pseudocolor, which were performed in each optical channel separately.

### RNA intereference (RNAi)

RNAi against Kcnn3 and Ezh2 was performed using Dharmacon ON-TARGETplus SMARTpool siRNA or corresponding scrambled sequences (25 nM) and DharmaFECT 1 Transfection Reagent (Thermo Fisher) according to the manufacturer’s protocol. Efficacy was assessed after 6 (Kcnn3) or 14 days (Ezh2) by Western blotting.

### Patch clamp

Pipettes were pulled from KG-12 glass (Kimble Garner) on a P97 puller (Sutter Instrument Co., Novato, CA) to a resistance of 2-5 MΩ and coated with HIPEC R6101 semiconductor protectant (Dow Corning, Auburn, MI). Current-clamp recordings were taken in gap-free mode on an Axopatch 200B amplifier with pClamp 10 software (Molecular Devices, Sunnyvale, CA). The extracellular solution contained (mM): 150 Na^+^, 160 Cl^−^, 5 K^+^, 2.5 Ca^2+^, 5 HEPES, and 5.5 glucose, pH 7.35, 300 mmol/kg osmolality. The intracellular solution contained (mM): 135 K^+^, 130 CH_3_SO_3_^−^, 20 Cl^−^, 5 Na^+^, 5 Mg^2+^, 5 HEPES, and 2 EGTA, pH 7.0, 290 mmol/kg osmolality in the pipette tip with 0.5 mM amphotericin B dissolved in DMSO at a final concentration of 0.8% in the backfill solution.

### Gene expression analysis by microarrays

Total RNA was extracted using the Norgen Biotek (Thorold, ON, Canada) Total RNA Purification Kit. RNA quality and quantity were tested using RNA 6000 Nano Kit (Agilent Technologies, Palo Alto, CA). Affymetrix (Santa Clara, CA) 3’ IVT Express Kit, GeneChip^®^ Mouse Genome 430 2.0 microarrays and GeneChip^®^ Scanner 3000 were used for expression profiling of ICL2A cells (n=3 independent cultures and microarrays) and control P6-9 C57BL/6J stomach *tunica muscularis* tissues (n=3 mice and microarrays). Data analysis was performed as previously described (Chen et al., 2007; Dave et al., 2015). The reproducibility and fidelity of the microarray method was tested by correlating gene-level expression data from biological replicates and with values obtained by qRT-PCR and mRNA sequencing (mRNA-sequencing; see next section) (**Table S2**). Following annotation using Affymetrix NetAffx 3’-IVT Expression Array annotation, release 33 (11/2/12), genes differentially expressed ((log_2_ fold change≥1 *OR* ≤−1) *AND* Benjamini-Hochberg *Q*≤0.05) between ICL2A cells and C57BL/6J unfractionated *tunica muscularis* were compared to corresponding data previously generated on the same platform in FACS-purified murine small intestinal myenteric and deep muscular plexus-associated ICC (ICC-MY and ICC-DMP, respectively (GEO accession number: GSE7809) (**Supplemental Datasets 1**). Ingenuity Pathway Analysis was used to identify significant (*P*<0.05) canonical pathways associated with gene subsets (**Supplemental Datasets 2**).

### Gene Expression Analysis by mRNA Sequencing

Total RNA was isolated and purified using the Qiagen (Valencia, CA) RNeasy Mini Kit. Libraries were constructed using the Illumina (San Diego, CA) TruSeq mRNA Sample Preparation Kit v2. Expression libraries were sequenced (paired-end; 51 bases/read) on the Illumina HiSeq 2000 platform. Data were analyzed using the MAP-RSeq pipeline. Reproducibility was verified by correlation analysis of biological replicates (n=2/cell line/condition) (**Table S2**).

### Chromatin immunoprecipitation-deep sequencing (ChIP-sequencing) and ChIP-qPCR

DNA-protein complexes in ICL2A and GIST-T1 cells (n=2 independent cultures) were crosslinked using fresh 1% formaldehyde (Thermo Fisher) for 10 min at room temperature followed by glycine quenching (125 mM, 5 min, room temperature). GIST-T1 cells were generously donated by Dr. Takahiro Taguchi, Division of Human Health and Medical Science, Graduate School of Kuroshio Science, Kochi University, Kochi, Japan) (Hayashi et al., 2015). These cells have not been authenticated or tested for Mycoplasma. Sorted genetically labeled ICC were subjected to cross-linking and quenching before FACS. Approximately 10^6^ cells were harvested and lysed by incubating in 10 mM Tris-HCl, pH 7.5, 10 mM NaCl, 0.5% NP-40 lysis buffer for 10 min at 4 °C. Chromatin was first fragmented using micrococcal nuclease at 2000 gel units/mL (New England Biolabs, Ipswich, MA) in 20 mM Tris-HCl, pH 7.5, 15 mM NaCl, 60 mM KCl, 1 mM CaCl_2_ digestion buffer at 37 °C. After 20 min, the reaction was stopped with 100 mM Tris-HCl, pH 8.1, 20 mM EDTA, 200 mM NaCl, 2% Triton X-100, 0.2% sodium deoxycholate buffer. The chromatin was further fragmented using a Bioruptor sonicator (Diagenode, Denville, NJ) at ‘high’ setting (15 cycles of 30 s on and 30 s off). Fragment sizes were determined using an Advanced Analytical Technologies (Ankeny, IA) Fragment Analyzer™ Automated CE System. Chromatin preparations were reacted with extensively validated (Landt et al., 2012) ChIP-grade antibodies against Ezh2, H3K27me3, H3K27ac, H3K4me3 and H3K9me3 (see **Supplemental Experimental Procedures** for antibody details; overnight at 4 °C) and precipitated using Protein-G agarose beads (Roche Diagnostics, Indianapolis, IN). After elution of DNA-protein complexes, crosslinks were reversed by proteinase K digestion overnight at 62 °C and DNA was purified using Qiagen PCR Purification Kit. Enrichment of target DNA relative to 1% of input DNA was analyzed by PCR against the transcription start site of the constitutively repressed gene *T* (brachyury) and the constitutively transcribed gene *Actb* (controls), as well as *Kit, Ano1*, and *Cacna1h* (see **Supplemental Experimental Procedures** for primer sequence).

Sequencing libraries were constructed using the NuGEN Ovation Ultralow DR Multiplex System 1-8 (NuGEN Technologies, San Carlos, CA). Briefly, the DNA fragments underwent end repair, barcoded sequencing adapter ligation, purification (Agencourt RNAClean XP beads; Beckman Coulter) and PCR-amplification (13 cycles). Final ChIP libraries were sequenced (paired-end; 51 bases/read) on the Illumina HiSeq 2000 platform. Data were analyzed by the HiChIP pipeline (Yan et al., 2014; Yan et al., 2016). Briefly, reads were aligned to the mm10 genome assembly using BWA and visualized using Integrative Genomics Browser (IGV(Robinson et al., 2011)). Mapped reads were post-processed to remove duplicates and pairs of reads mapping to multiple locations and tag density files were generated for all genes in the Affymetrix NetAffx 3’-IVT Expression Array annotation, release 33 (11/2/12) within 5 kb of TSS’s. *MACS2* (Zhang et al., 2008) (for H3K27ac and H3K4me3) and *SICER* (Zang et al., 2009) (for H3K27me3 and H3K9me3) algorithms were used for peak-calling in relation to the input DNA. Super-enhancers were detected with Rank Ordering of Super-Enhancers (*ROSE*) algorithm (http://younglab.wi.mit.edu/super_enhancer_code.html) (Loven et al., 2013; Whyte et al., 2013). Reproducibility was verified by correlation analysis of biological replicates (n=2/cell line/condition) (**Table S2**). H3K27ac data from sorted small intestinal ICC was correlated with data from colonic ICC. Data were visualized using *pheatmap* in R (https://www.r-project.org/).

### Assessment of membrane potential by flow cytometry using DiBAC_4_(3)

Membrane potential of ICL2A cells was determined by flow cytometry using the voltage-sensitive fluorescent dye DiBAC_4_(3) (Thermo Fisher) using methods slightly modified from a published protocol (Klapperstuck et al., 2009). Briefly, cells were lifted by trypsinization, re-suspended to 400,000 cells/mL Ca^2+^ - and Mg^2+^-free Hank’s Balanced Salt Solution containing 5% fetal bovine serum supplemented with 0 or 2 mM CaCl_2_ and incubated with 50 nM DiBAC_4_(3) at 37 °C. After 35 min, apamin (Sigma-Aldrich), a highly selective inhibitor of Kcnn1, Kcnn2 and Kcnn3 channels (Adelman et al., 2012), or vehicle (H_2_O) was added to the suspensions (final concentration: 300 nM). After 5 min, nonviable cells were labeled with 5 μg/mL propidium iodide (BioSure) at room temperature for 5 min. Background-subtracted median DiBAC_4_(3) fluorescence values (F) were obtained using a Becton Dickinson LSR II flow cytometer (see configuration in **Supplemental Experimental Procedures**) and FlowJo 10 (Tree Star). Cells treated with 1-2 μg/mL gramicidin A (Sigma-Aldrich) for 40 min at 37 °C were used to determine the maximum achievable level of DiBAC_4_(3) fluorescence corresponding to depolarization to 0 mV (F_dep_). Membrane potential was calculated using the formula V_m_= −61*log(F_dep_/F).

### Drugs not mentioned elsewhere

Adenosine dialdehyde (Adox) and BIX-0129 were from Sigma-Aldrich, EPZ-6438 and GSK126 were from Xcessbio Biosciences, Inc.(San Diego, CA) suberanilohydroxamic acid (SAHA) was from Santa Cruz Biotechnology, Inc. (Dallas, TX) and RG108 was from Cayman Chemical Co.(Ann Arbor, MI).

### Statistical analyses

Mice were randomly assigned to various experimental groups. Data were analyzed by nonparametric methods including Mann-Whitney rank sum test, Wilcoxon signed rank test, Kruskal-Wallis one-way analysis of variance (ANOVA) on ranks and Spearman’s rank order correlation test. *P*<0.05 was considered significant. In figure descriptions, unless otherwise specified, Mann-Whitney rank sum test was used.

### Accession Numbers

Microarray, mRNA-sequencing and ChIP-sequencing data have been deposited in the National Center for Biotechnology Information Gene Expression Omnibus (http://www.ncbi.nlm.nih.gov/geo/) under SuperSeries accession number GSE77838.

## AUTHOR CONTRIBUTIONS

T.O. supervised the project and developed the overall study hypothesis. T.O., S.A.S., Y.H., G.B.G. and S.M. designed the experiments. Y.H., S.A.S., J.H.L., A.L., P.R.S., G.B.G., S.M., J.J.R., S.J.G., V.J.H., M.R.B. and D.D.R. performed experiments and data analysis. H.Y., J.N. and S.A.S. performed bioinformatic analysis. S.K., D.S. and R.A.U. provided critical research resources and consultation on the use of animal models. T.O., R.A.U., Z.Z. and G.F. provided guidance on data interpretation and important feedback. S.A.S. drafted the manuscript. All authors have read and approved the final manuscript. S.A.S. and Y.H. contributed equally to this study.

## Supporting information

## ACKNOWLEDGMENTS

We thank G.J. Gores (Division of Gastroenterology and Hepatology, Mayo Clinic, Rochester, MN) for granting us access to the LI-COR Odyssey Scanner. We also thank the Mayo Clinic Microscopy and Cell Analysis Core (director: Jeffrey L. Salisbury) for fluorescence-activated cell sorting and the Medical Genome Facility (director: Eric D. Wieben) for the high-throughput sequencing and the Affymetrix microarray analysis. Supported in part by US National Institutes of Health grants R01DK058185, P01DK068055 and P30DK084567, The Life Raft Group (https://liferaftgroup.org/) and the Mayo Clinic Center for Individualized Medicine (http://mayoresearch.mayo.edu/center-for-individualized-medicine).

## COMPETING FINANCIAL INTERESTS

The authors declare no competing financial interests.

