## Supplementary material for "Ezh2-dependent epigenetic reprogramming controls a developmental switch between modes of gastric neuromuscular regulation"

### SUPPLEMENTAL FIGURES AND SUPPLEMENTAL TABLES

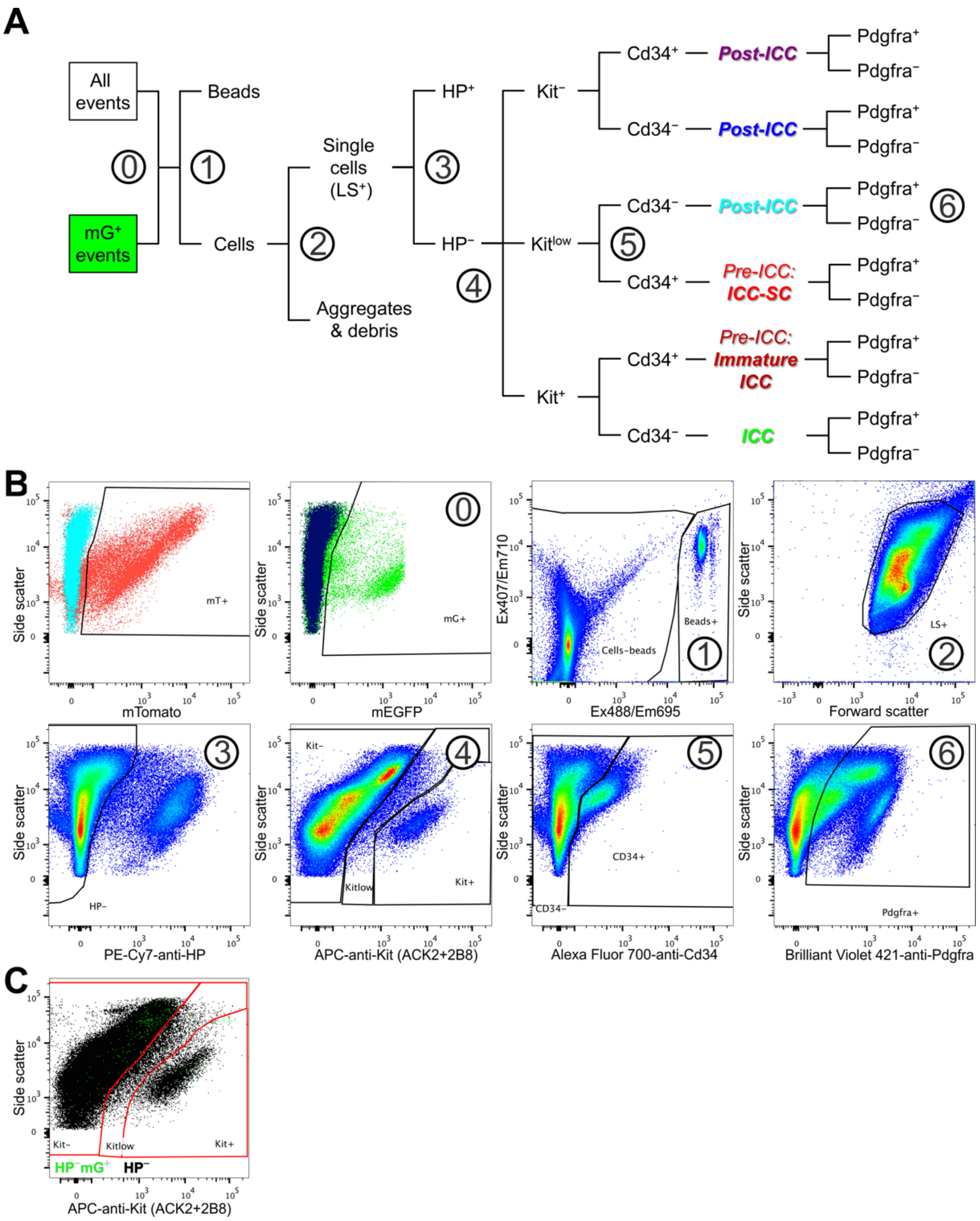


**Figure S1** Flow cytometry analysis of genetically labeled cells of the ICC lineage in *Kit*$^{CreERT2/+}$;*R26*$^{mT\text{-}mG/mT\text{-}mG}$ mice, Related to Figure 1. (**A**) Gating sequence and identification of members of the ICC lineage. ICC, immature ICC and ICC-SC were detected in the HP$^{-}$ subset as Kit$^{+}$Cd34$^{-}$, Kit$^{+}$Cd34$^{+}$ and Kit$^{low}$Cd34$^{+}$ cells, respectively(Bardsley et al., 2010; Lorincz et al., 2008), following the exclusion of counting beads, cell aggregates and debris. Anticipated phenotypes of post-ICC populations were defined based on refs.(Lorincz et al., 2008; Torihashi et al., 1999) (Kit$^{low/-}$Cd34$^{-}$) and ref.(Iino and Nojyo, 2009) (Kit$^{-}$Cd34$^{+}$). (**B**) Representative projections illustrating key cell populations and decision forks identified by numbers in **A**. Population boundaries were drawn along lines representing identical cell counts in quantile contour plots. First panel: detection of cells expressing membrane-targeted Tomato (mTomato, mT) from the non-recombined *R26*$^{mT\text{-}mG/mT\text{-}mG}$ reporter gene. ⓪: Identification of cells expressing membrane-targeted enhanced green fluorescent protein (mEGFP, mG). ①: Detection of counting beads. Ex, excitation wavelength and Em, emission maximum for the fluorescent channels used. ②: Cells with light scatter properties characteristic of single live cells (LS$^{+}$). ③: Separation of HP$^{-}$ cells from cells of hematopoietic origin (Cd45$^{+}$Cd11b$^{+}$F4/80$^{+}$ leukocytes). PE-Cy7, phycoerythrin-cyanine 7 tandem conjugate. ④: Detection of Kit$^{-}$, Kit$^{low}$ and Kit$^{+}$ cells. APC, allophycocyanin. ⑤: Separation of Cd34$^{-}$ and Cd34$^{+}$ cells. ⑥: Identification of Pdgfra$^{+}$ and Pdgfra$^{-}$ cells. (**C**) Nonspecific mG fluorescence (green) in a vehicle-treated *Kit*$^{CreERT2/+}$*;R26*$^{mT\text{-}mG/mT\text{-}mG}$ mouse. Dot plot of 100,000 HP$^{-}$ cells (black) is shown.

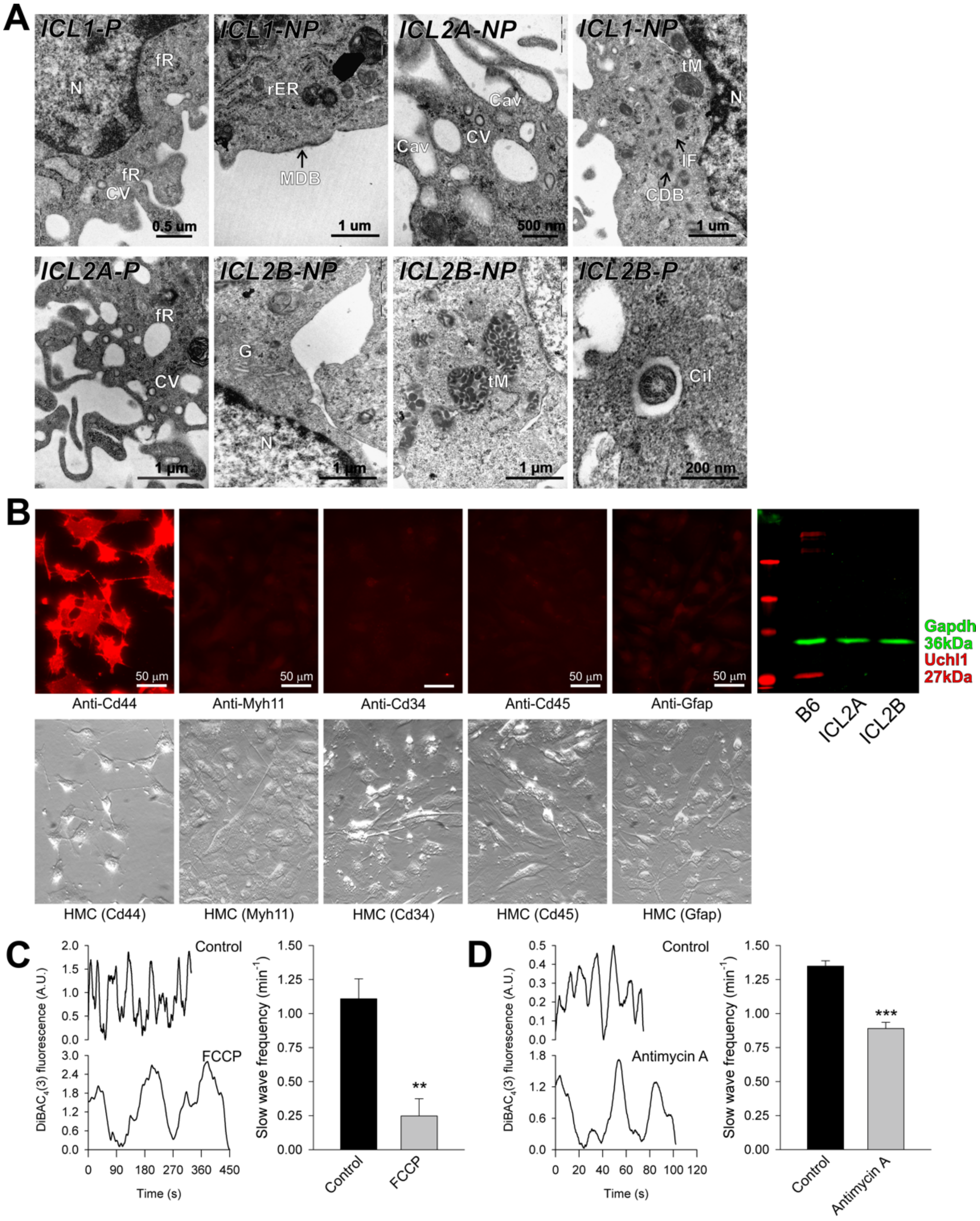


**Figure S2** Conditionally immortalized ICC retain key ICC traits in early/intermediate passages, Related to Figure 2. (**A**) Ultrastructural features considered hallmarks of ICC. P and NP indicate cells maintained under conditions permissive and nonpermissive, respectively, for the tsTAg. N, nucleus; CV, coated vesicle; fR, free ribosomes; rER, rough endoplasmic reticulum; MDB, membrane-associated dense band; Cav, caveolae; CDB, cytoplasmic dense band; IF, intermediate filaments; tM, mitochondria with tubular cristae; G, Golgi zones; Cil, primary cilium (9×2+0). (**B**) Conditionally immortalized, clonally derived ICC express the ICC marker Cd44 but not the smooth muscle marker Myh11, the ICC-SC marker Cd34, the hematopoietic marker Cd45 or the neural crest markers Gfap and Uchl1. Representative immunofluorescent images from ICL2A cultures (n=2; passage 11; nonpermissive conditions) and a Western blot from ICL2A (passage 14) and ICL2B (passage 16) cultures maintained under semipermissive conditions are shown (n=6 cultures/cell line). B6, corpus+antrum muscles from C57BL6/J mice (n=6). (**C**) Inhibition of phasic electrical activity in ICL2B cells by 30-min exposure to carbonilcyanide p-triflouromethoxyphenylhydrazone (FCCP, 1 μM), an uncoupler of mitochondrial oxidative phorsphorylation (means±s.e.m.; Wilcoxon signed rank test, **, *P*=0.004, *W*=-45.000, n=8). (**D**) Inhibition of phasic electrical activity in ICL1 (n=10) and ICL2B cells (n=5) by antimycin A (10 μM), an inhibitor of mitochondrial electron transport chain complex III (means±s.e.m.; Wilcoxon signed rank test, ***, *P*<0.001, *W*=-120.000). Both FCCP and antimycin completely abolished $DiBAC_4(3)$ oscillations after 1 h exposure (not shown). $DiBAC_4(3)$ imaging was performed in early-passage (<8) cells maintained under nonpermissive conditions for the tsTAg. A.U., arbitrary units.

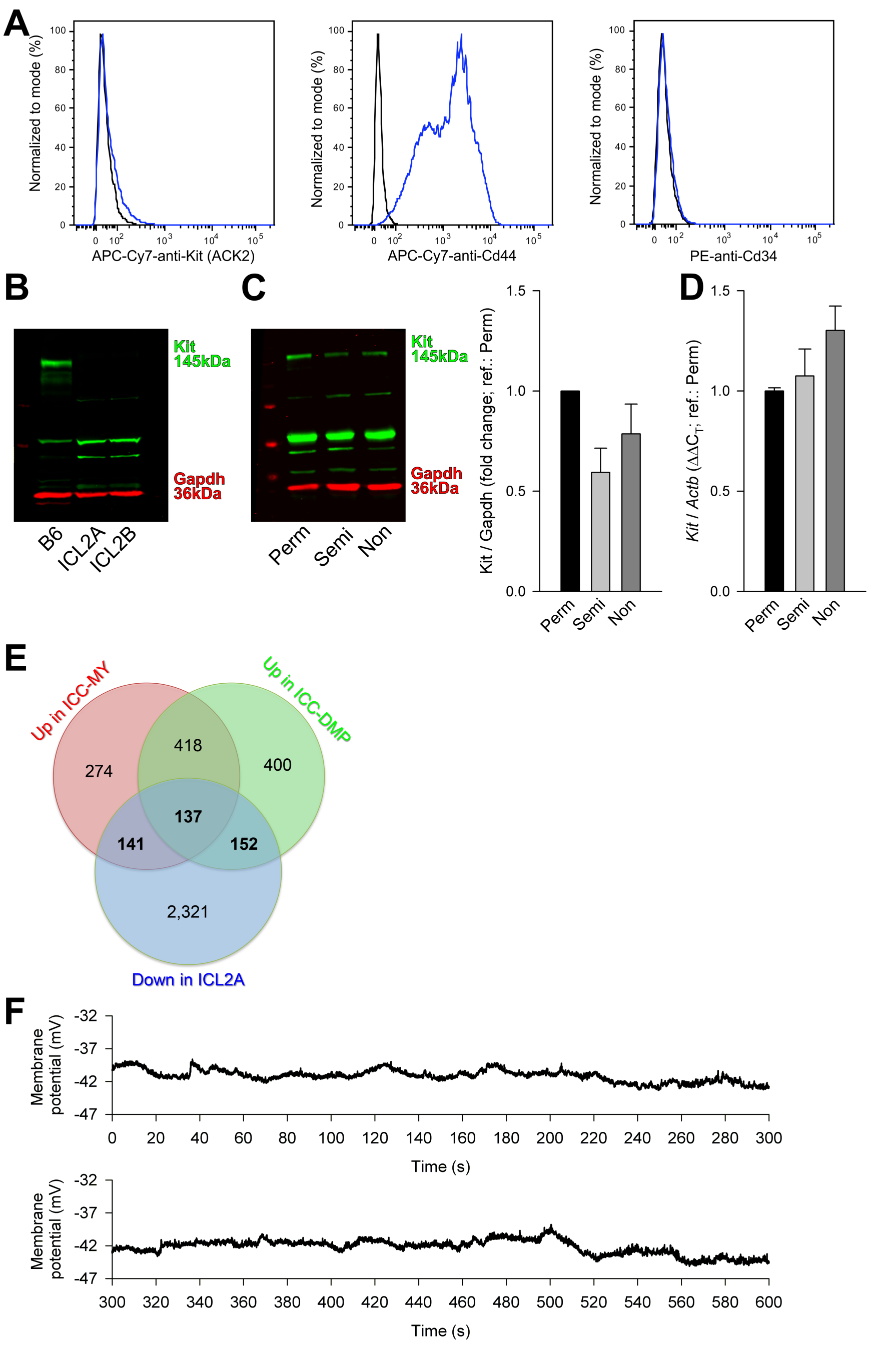


**Figure S3** Conditionally immortalized ICC lose ICC phenotype with extended culturing, Related to Figure 3. (**A**) Analysis of Kit, Cd44 and Cd34 expression by flow cytometry in late-passage (31) ICL2B cells maintained under semipermissive conditions. Representatives of 3 experiments are shown. (**B**) Reduced Kit expression detected by Western blotting in ICL2A (passage 20) and ICL2B cells (passage 24) vs. C57BL/6J (B6) corpus+antrum *tunica muscularis*. Uncropped version of the Western blot image in **Figure 3C** is shown (a representative of 4 experiments). (**D**,**D**) Kit protein (**C**) and mRNA (**D**) expression (means±s.e.m.) in ICL2A cells is not significantly affected by culturing under permissive (Perm), semipermissive (Semi) or nonpermissive (Non) conditions for 4 days (Western blots: $P$=0.127, $H$(2)= 4.127, n=5/condition; qRT-PCR: $P$=0.254, $H$(2)=3.200, n=3/condition; Kruskal-Wallis ANOVA on ranks). Note that overall low Kit expression necessitated the loading of relatively high amounts of protein and this resulted in strong nonspecific bands in the uncropped representative Western blot image. (**E**) Venn diagram showing subsets of genes significantly overexpressed in ICC-MY and ICC-DMP relative to their source tissue and significantly downregulated in late-passage ICL2A cells maintained under semipermissive conditions. Significant differential expression was defined as $\log_2$ fold change≤-1 or ≥1 with Benjamini-Hochberg $Q$<0.05. Numbers in bold indicate gene sets analyzed by IPA (**Supplemental Dataset 2**). (**F**) A representative patch clamp (current clamp) trace demonstrating lack of phasic activity in an extended recording performed in an ICL2B cell.

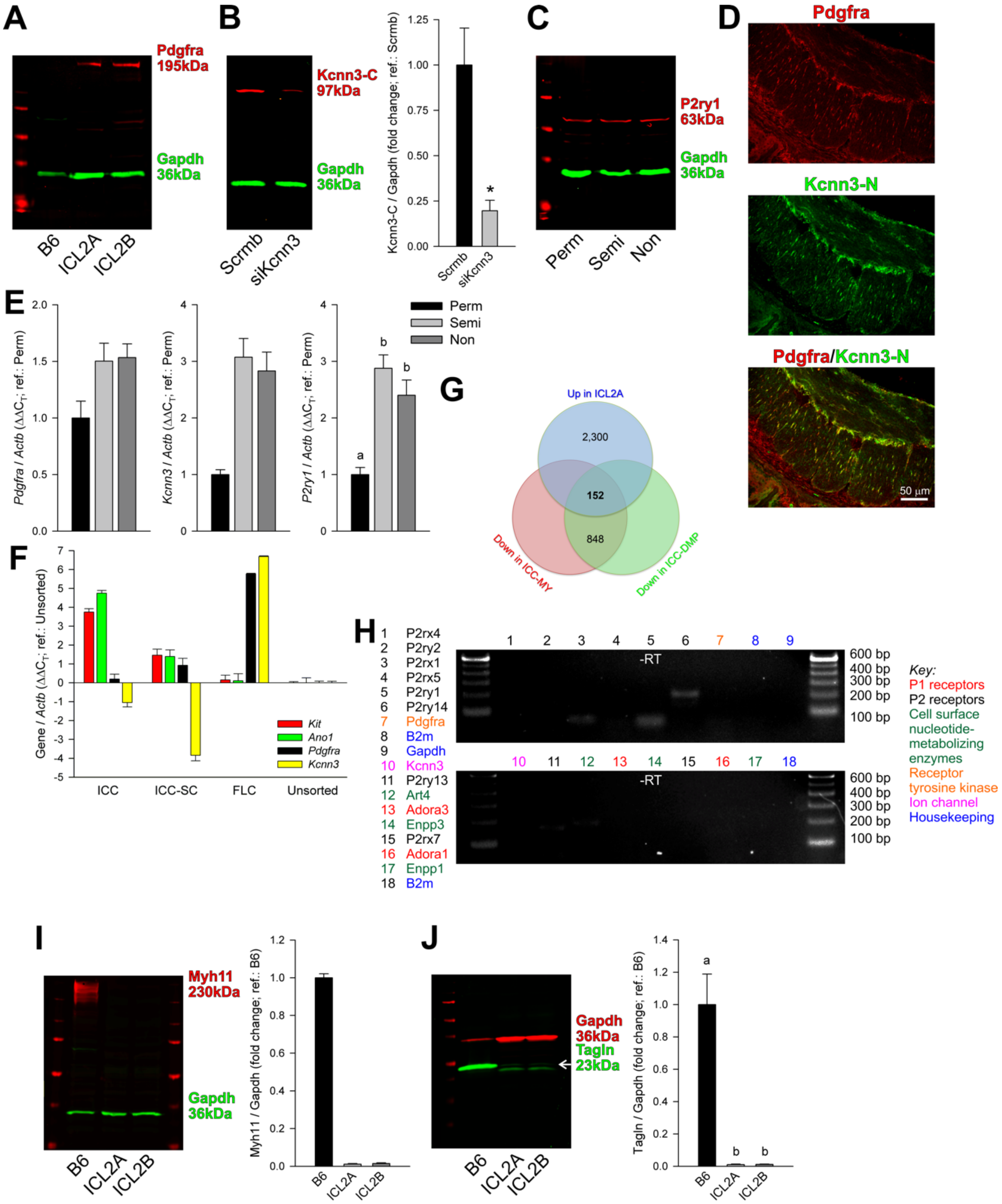

**Figure S4** Conditionally immortalized $Kit^{low/-}$ ICC gain FLC but not smooth muscle traits, Related to Figure 4. (**A**) Pdgfra expression detected by Western blotting in passage 18 ICL2A and passage 20 ICL2B cells vs. C57BL/6J (B6) corpus+antrum muscles. Uncropped version of the Western blot image in **Figure 4C** is shown (representative of 3 experiments). (**B**) Demonstration of the specificity of the Kcnn3-C antibody by RNAi-mediated knockdown of Kcnn3 (siKcnn3; 6 days) in ICL2A cells (nonpermissive). Scrmb, scrambled siRNA. A representative full blot and quantitative data from 4 experiments are shown ($P$=0.029). (**C**) P2ry1 expression in ICL2A cells cultured under permissive (Perm), semipermissive (Semi) or nonpermissive (Non) conditions for 4 days. A representative full blot from 4 experiments is shown. (**D**) Kcnn3-N antibody detects Kcnn3 (green) exclusively in $Pdgfra^{+}$ cells (red) of the stomach of nondiabetic NOD/ShiLtJ mice. Yellow color signifies co-localization. Representative images from 12 sections of 4 mouse stomachs are shown. (**E**) Detection of *Pdgfra*, *Kcnn3* and *P2ry1* mRNA by qRT-PCR (means±s.e.m.; n=3/group) in ICL2A cells cultured under permissive (Perm), semipermissive (Semi) or nonpermissive (Non) conditions for 4 days. Groups not sharing the same superscript are different by Student-Newman-Keuls test ($P$<0.05) following Kruskal-Wallis nonparametric ANOVA (*Pdgfra*: $P$=0.132, $H$(2)=4.356; *Kcnn3*: $P$=0.05, $H$(2)=5.793; *P2ry1*: $P$=0.025, $H$(2)=5.956). (**F**) Validation of *Pdgfra* and *Kcnn3* qRT-PCR primers used in experiments depicted in panel **E**. Gastric ICC, ICC-SC and FLC were FACS-purified from BALB/c mouse stomachs as $HP^{-}Kit^{+}Cd44^{+}Cd34^{-}$, $HP^{-}Kit^{low}Cd44^{+}Cd34^{+}$ and $HP^{-}Kit^{-}Cd44^{-}Cd34^{-}Pdgfra^{+}$ cells, respectively, as described previously(Schwamb et al., 2015). (**G**) Venn diagram showing subsets of genes significantly downregulated in ICC-MY and ICC-DMP relative to their source tissue and significantly upregulated in late-passage ICL2A cells (semipermissive). Significant differential expression was defined as $\log_2$ fold change≤−1 or ≥1 with Benjamini-Hochberg $Q$<0.05. Number in bold indicates gene set analyzed by IPA (**Supplemental Datasets 2**). (**H**) Detection of FLC-specific genes by RT-PCR using primers from ref.(Peri et al., 2013). Representative data from 3 experiments are shown.(**I,J**) Passage 20 ICL2A and passage 24 ICL2B cells (semipermissive) do not express the smooth muscle markers Myh11 (**I**) and Tagln (also known as SM-22; (**J**). Representative immunoblots and quantitative data vs. C57BL/6J (B6) corpus+antrum muscles (means±s.e.m.) are shown. Groups not sharing the same superscript are different by Student-Newman-Keuls test ($P$<0.05) following Kruskal-Wallis nonparametric ANOVA (Myh11: $P$=0.05, $H$(2)=5.600, n=3/group; Tagln: $P$=0.009, $H$(2)=9.380, n=5/group).

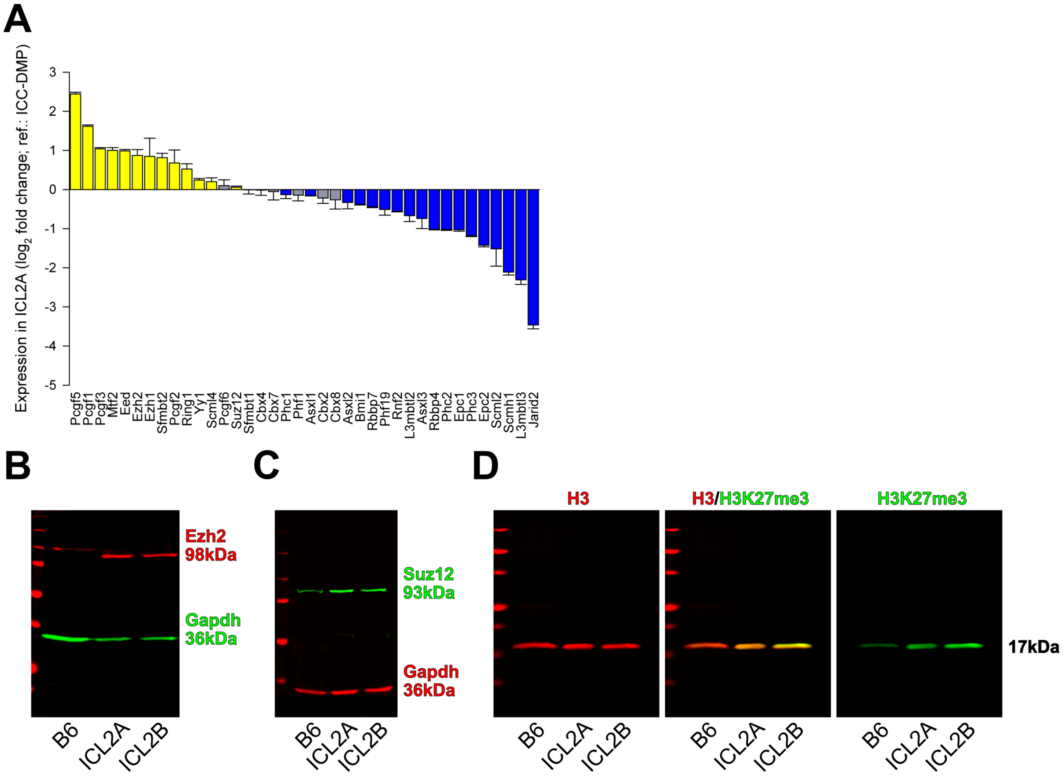


**Figure S5** H3K27me3 occupies ICC genes after phenotypic transition into FLC, Related to Figure 5. (**A**) Expression of PcG genes(Sauvageau and Sauvageau, 2010; Woo et al., 2010) in ICL2A cells relative to ICC-DMP cells detected by microarray analysis (n=3/group; semipermissive). Yellow and blue indicate genes up- and downregulated in all ICL2A samples; gray: inconsistent change among replicates. (**B**-**D**) Detection of Ezh2, Suz12 and H3K27me3 by Western blotting in passage 20 ICL2A and passage 24 ICL2B cells (semipermissive) vs. C57BL/6J (B6) corpus+antrum muscles. Uncropped versions of the Western blot images in **Figure 5B** are shown. Blots are representatives of 5 (Ezh2, Suz12) and 7 experiments (H3K27me3).

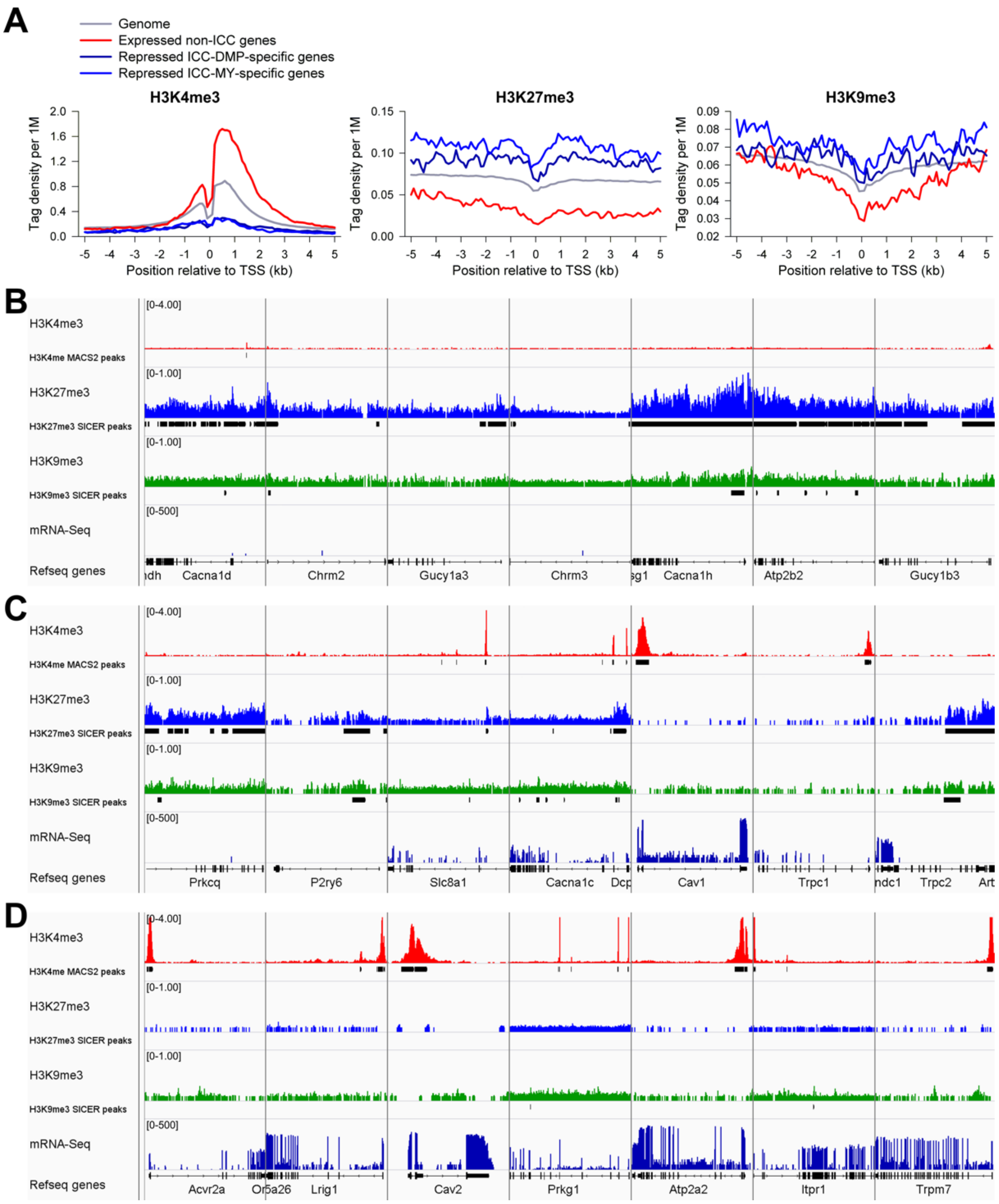


**Figure S6** H3K27me3 occupies ICC genes after phenotypic transition into FLC, Related to Figure 5. (**A**-**D**) ChIP-sequencing on H3K4me3, H3K27me3 and H3K9me3 and mRNA-sequencing results obtained in ICL2A cells are shown. All displayed data are representatives of 2 independent experiments. (**A**) Tag density plots (TSS±5 kb) from ChIP-sequencing averaged for the indicated gene sets or the entire genome (gray). Note inverse correlation between transcription and H3K27me3 occupancy and direct correlation between transcription and H3K4me3 occupancy. H3K9me3 occupancy was less correlated with gene transcription. (**B**-**D**) Genome browser tracks from the ChIP-sequencing (values: RPM) and mRNA-sequencing (values: log RPKM) experiments. ICC genes fell into 3 groups: not expressed and occupied by H3K27me3 (with or without H3K9me3 binding: *Cacna1d*, *Chrm2*, *Gucy1a3*, *Chrm3*, *Cacna1h*, *Atp2b2*, *Gucy1b3*, *Prkcq*, *P2ry6* and *Trpc2*), lowly expressed and bearing both H3K4me3 and H3K27me3 peaks (*Slc8a1* and *Cacna1c*) and expressed and occupied by H3K4me3 only (*Trpc1*, *Acvr2a*, *Lrig1*, *Cav2*, *Prkg1*, *Atp2a2*, *Itpr1* and *Trpm7*).

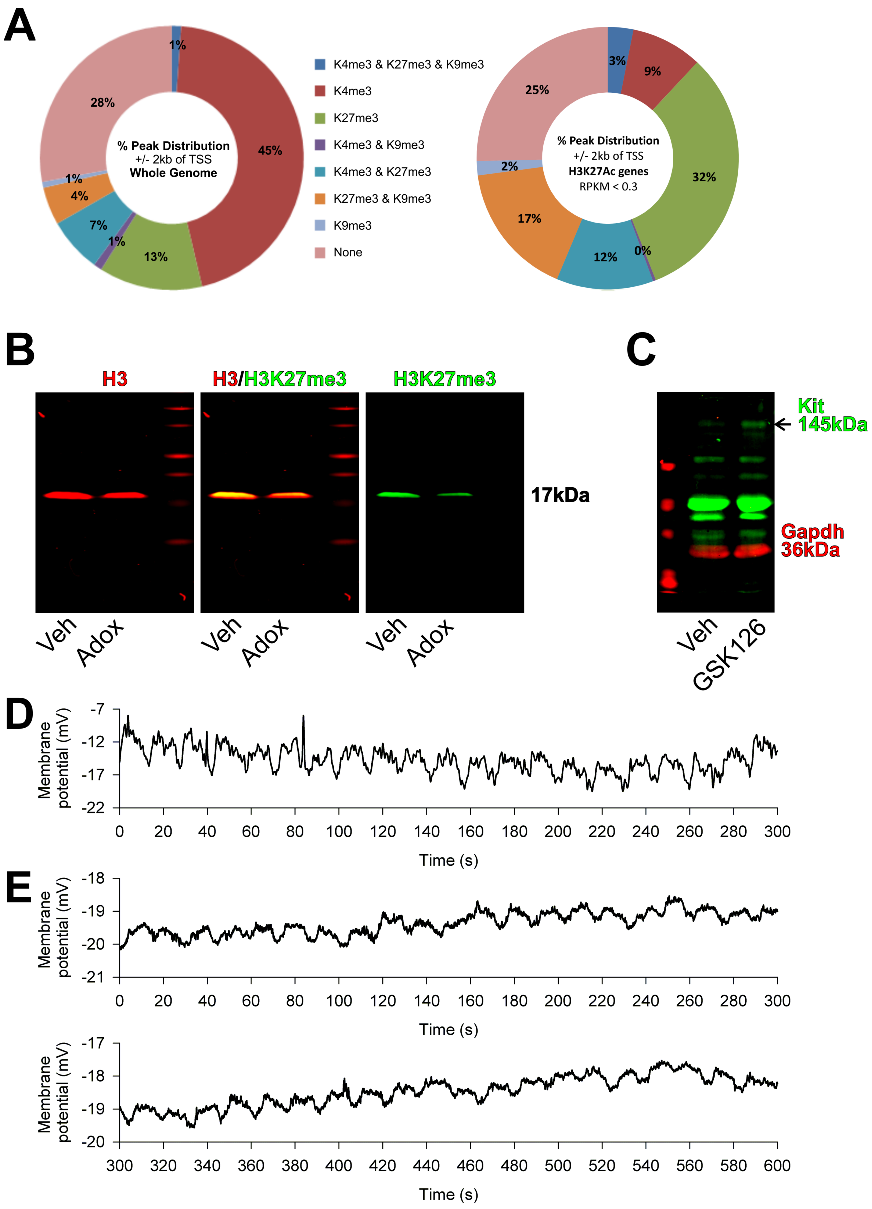


**Figure S7** Ezh2 inhibition restores ICC phenotype and function, Related to Figure 7. (**A**) Left: Pie chart showing the percentage of significant histone mark peaks and their combinations within TSS±2 kb of all genes in ICL2A cells (semipermissive). Right: corresponding data for H3K27ac-marked ICC genes with low expression in ICL2A cells (RPKM<0.3(Ramskold et al., 2009); **Supplemental Datasets 1**). Note lower frequency of genes occupied by H3K4me3 and higher frequency of genes bearing H3K27me3 (and, to a lesser extent, H3K9me3) in the subset repressed in ICL2A cells but bearing H3K27ac marks in ICC. (**B**) In passage 18 ICL2A cells (permissive), 2.5 μM Adox for 4 days reduced H3K27me3 levels. A representative Western blot from 5 experiments is shown. Veh, vehicle. (**C**) In passage 22 ICL2A cells (permissive), 2-week treatment with 500 nM GSK126 increased Kit protein expression. A representative Western blot from 6 experiments is shown. Veh, vehicle. (**D-E**) Extended traces from the recordings illustrated in **Figure 7F** are shown.

**Table S1** Transmission electron microscopic characteristics of conditionally immortalized interstitial cell of Cajal (ICC) lines, Related to Figure 2.

| **Organelles** | **ICL1 P**[a] | **ICL1 NP**[b] | **ICL2A P** | **ICL2A NP** | **ICL2B P** | **ICL2B NP** |
|---|---|---|---|---|---|---|
| Caveolae | + | ++ | + | ++ | + | ++ |
| Membrane-associated dense bands | + | + | - | - | - | + |
| Basal lamina | (+) | (+) | - | - | - | (+) |
| Cytoplasmic dense bodies | + | + | + | - | - | - |
| Rough endoplasmic reticulum | + | ++ | ++ | ++ | + | ++ |
| Smooth endoplasmic reticulum | ++ | + | - | ++ | + | + |
| Golgi apparatus | + | - | - | ++ | + | ++ |
| Mitochondria | ++ | +++ | ++ | +++ | + | ++ |
| Coated vesicles | ++ | + | ++ | + | + | + |
| Free ribosomes | ++ | ++ | ++ | ++ | ++ | ++ |
| Intermediate (10 nm) filaments | ++ | ++ | ++ | ++ | + | + |
| Mitoses | - | - | + | + | + | + |
| Centrioles | + | + | - | - | + | + |
| Cilia | (+) | (+) | - | - | + | + |

[a], P, cultured under permissive conditions for the SV40 tsA58 mutant large T antigen; [b], NP, cultured under nonpermissive conditions for the SV40 tsA58 mutant large T antigen

**Table S2** Gene expression microarray, mRNA-sequencing (mRNA-seq) and chromatin immunoprecipitation-sequencing (ChIP-seq) correlation analysis results, Related to Figure 3,5,6.

| Cell line | Analysis method(s) | Variable 1 | Variable 2 | ρ[a] | n | *P* |
|---|---|---|---|---|---|---|
| ICL2A | Microarray[b] (gene-level[c]) | $log_2$ MAS5 expression; SP[d] replicate 1 | $log_2$ MAS5 expression; SP replicate 1 | 0.948 | 21524 | <0.001 |
| ICL2A | Microarray (gene-level) | $log_2$ MAS5 expression; SP replicate 3 | $log_2$ MAS5 expression; SP replicate 2 | 0.947 | 21524 | <0.001 |
| ICL2A | Microarray (gene-level) | $log_2$ MAS5 expression; SP replicate 1 | $log_2$ MAS5 expression; SP replicate 3 | 0.95 | 21524 | <0.001 |
| ICL2A | mRNA-seq | $log_2$ RPKM[e]; P[f] replicate 1 | $log_2$ RPKM; P replicate 2 | 0.994 | 15275[g] | <0.001 |
| ICL2B | mRNA-seq | $log_2$ RPKM; P replicate 1 | $log_2$ RPKM; P replicate 2 | 0.993 | 15255[g] | <0.001 |
| ICL2A | mRNA-seq | $log_2$ RPKM; NP[h] replicate 1 | $log_2$ RPKM; NP replicate 2 | 0.922 | 15100[g] | <0.001 |
| ICL2B | mRNA-seq | $log_2$ RPKM; NP replicate 1 | $log_2$ RPKM; NP replicate 2 | 0.935 | 15069[g] | <0.001 |
| ICL2A | mRNA-seq | $log_2$ RPKM; P average of 2 | $log_2$ RPKM; NP average of 2 | 0.932 | 16034[g] | <0.001 |
| ICL2A | mRNA-seq, microarray (gene-level) | $log_2$ RPKM; P average of 2 | $log_2$ MAS5 expression; SP average of 3 | 0.817 | 12709[i] | <0.001 |
| ICL2A | mRNA-seq, microarray (gene-level) | $log_2$ RPKM; NP average of 2 | $log_2$ MAS5 expression; SP average of 3 | 0.87 | 13260[i] | <0.001 |
| ICL2A | Microarray (gene-level[j]), qRT-PCR[k] | Microarray $log_2$ fold change (SP) vs. source tissue, average of 3 | qRT-PCR $log_2$ fold change (SP) vs. source tissue, average of 3 | 0.769 | 50 | <0.001 |
| ICL2A | ChIP-seq: H3K4me3 | $log_2$ NTC[l]; NP replicate 1 | $log_2$ NTC; NP replicate 2 | 0.953 | 41695[m] | <0.001 |
| ICL2A | ChIP-seq: H3K27me3 | $log_2$ NTC; NP replicate 1 | $log_2$ NTC; NP replicate 2 | 0.982 | 29992[m] | <0.001 |
| ICL2A | ChIP-seq: H3K9me3 | $log_2$ NTC; NP replicate 1 | $log_2$ NTC; NP replicate 2 | 0.925 | 22930[m] | <0.001 |
| GIST-T1 | ChIP-seq: H3K4me3 | $log_2$ NTC; replicate 1 | $log_2$ NTC; replicate 2 | 0.936 | 46666[m] | <0.001 |
| GIST-T1 | ChIP-seq: H3K27me3 | $log_2$ NTC; replicate 1 | $log_2$ NTC; replicate 2 | 0.908 | 30462[m] | <0.001 |
| ICC[n] | ChIP-seq: H3K27ac | $log_2$ NTC; Small intestine | $log_2$ NTC; Colon | 0.658 | 73157[m] | <0.001 |

[a], Spearman rank order correlation coefficient; [b], Affymetrix (Santa Clara, CA) Mouse Genome 430.2; [c], For genes represented by more than one probe set, the probe set with the lowest MAS5 detection *P* value was used. In case of ties, the probe set with the highest MAS5 expression value was used; [d], SP, semipermissive culture conditions; [e], RPKM, reads per kilobase of transcript per million mapped reads; [f], P, permissive culture conditions; [g], Genes with

non-zero RPKM; [h], NP, nonpermissive culture conditions; [i], Genes with RPKM>0.047(Zhao et al., 2014); [j], For genes represented by more than one probe set, the probe set with the lowest Benjamini-Hochberg *Q* value (from differential gene expression analysis by the empirical Bayes approach following pre-processing with robust multiple-array analysis, RMA) was used; [k], qRT-PCR, real-time reverse transcription-polymerase chain reaction; [l], NTC, normalized tag count; [m], $\log_2$ NTC values>=2; [n], Interstitial cells of Cajal purified by fluorescence-activated cell sorting. See reference (Zhao et al., 2014).

**Supplemental Datasets 1 (Gene Lists).**

ICC-MY-specific genes. Gene set specific for ICC-MY (up [$\log_2$ fold change>1 *AND* Benjamini-Hochberg $Q<0.05$] in ICC-MY vs. source tissue *AND* not up [$\log_2$ fold change<1] in ICC-DMP vs. source tissue) and differential expression of this gene set in ICL2A cells vs. gastric muscles.

ICC-DMP-specific genes. Gene set specific for ICC-DMP (up in ICC-DMP vs. source tissue *AND* not up in ICC-MY vs. source tissue) and differential expression of this gene set in ICL2A cells vs. gastric muscles.

Genes common to all ICC. Gene set specific to both ICC-MY and ICC-DMP (up in ICC-MY vs. source tissue *AND* up in ICC-DMP vs. source tissue) and differential expression of this gene set in ICL2A cells vs. gastric muscles.

Non-ICC genes. Gene set downregulated in both ICC-MY and ICC-DMP (down [$\log_2$ fold change<−1 *AND* Benjamini-Hochberg $Q<0.05$] in ICC-MY vs. source tissue *AND* down in ICC-DMP vs. source tissue] and differential expression of this gene set in ICL2A cells vs. gastric muscles.

ICC-related genes (ICC-MY-specific genes *OR* ICC-DMP-specific genes *OR* genes common to all ICC) that are PcG targets and differential expression of this gene set in ICL2A cells vs. gastric muscles.

**Supplemental Datasets 2 (Ingenuity Pathway Analysis)**

ICC-MY-specific genes.

ICC-DMP-specific genes.

Genes common to all ICC.

Non-ICC genes significantly upregulated in ICL2A.

### SUPPLEMENTAL EXPERIMENTAL PROCEDURES

**Validation of *in-vivo* lineage tracing.** Cells with recombined reporter alleles (mG$^{+}$ cells) were enumerated in cell populations lacking hematopoietic markers (HP$^{–}$; to exclude Kit$^{+}$ mast cells) and expressing combinations of Kit, Cd34 and Pdgfra on their surface by flow cytometry in the whole gastric corpus+antrum *tunica muscularis* using established approaches (Bardsley et al., 2010) with modifications (see **Figures S1A,S1B** and next section). In *R26*$^{mT-mG}$ mice (n=16), the reporter construct was expressed in 89±3% (mean±s.e.m.) of HP$^{–}$ cells (**Figure S1B**). In control mice (n=10) including tamoxifen-treated *Kit*$^{+/+}$*;R26*$^{mT-mG/mT-mG}$ mice and untreated *Kit*$^{CreERT2/+}$*;R26*$^{mT-mG/mT-mG}$ and strain-matched wild-type mice, fluorescent cells in the channel used for the detection of mG did not exceed 0.93%, 0.52% and 0.11% in the HP$^{–}$Kit$^{+}$, HP$^{–}$Kit$^{low}$ and HP$^{–}$Kit$^{–}$ fractions, respectively (background; **Figure S1C**). This background fluorescence represented 1.16±0.06% of HP$^{–}$Kit$^{+}$mG$^{+}$ cells and 3.95±1.02% for HP$^{–}$Kit$^{low}$mG$^{+}$ cells in the experimental cohort (n=12). Recombination efficiency at P11 strongly correlated with median Kit expression across all samples and Kit gates (Kit$^{+}$, Kit$^{low}$ and Kit$^{–}$; Spearman's ρ=0.805, $P$<0.001, n=21). Reflecting this strong relationship between recombination and *Kit* transcription, in tamoxifen-treated, P11 *Kit*$^{CreERT2/+}$*;R26*$^{mT-mG/mT-mG}$ mice (n=7), background-subtracted mG$^{+}$ cell frequencies were 75±4% (HP$^{–}$Kit$^{+}$), 11±2% (HP$^{–}$Kit$^{low}$) and 0.31±0.06% (HP$^{–}$Kit$^{–}$) (mean±s.e.m.). Taking into account the average expression of the reporter construct (89±3% of the HP$^{–}$ cells; see above), the corrected recombination efficiencies were 84±5% (HP$^{–}$Kit$^{+}$), 13±2% (HP$^{–}$Kit$^{low}$) and 0.35±0.07% (HP$^{–}$Kit$^{–}$).

**Multi-parameter flow cytometry analysis of cells of the interstitial cell of Cajal (ICC) lineage.** ICC and related cells (see **Figures S1A,S1B**) were identified in the hematopoietic marker-negative fraction of dissociated mouse gastric corpus+antrum *tunica muscularis* using previously published protocols (Bardsley et al., 2010; Hayashi et al., 2015; Izbeki et al., 2010) with modifications (see below for antibodies and **Figures S1A,S1B** for gating scheme). Briefly, intact gastric corpus+antrum muscles were incubated with allophycocyanin (APC)-anti-Kit antibody (clone: ACK2) at 4 °C for 3 h, then dissociated with collagenase digestion, triturated and filtered. Single-cell suspensions were labeled with anti-mouse Cd16/32 antibody (Fcgr3/Fcgr2b; Fc block). Hematopoietic cells were identified with phycoerythrin (PE)-cyanine (Cy) 7-coupled anti-mouse Cd11b (Itgam), anti-Cd45 (Ptprc) and anti-F4/80 (Adgre1) antibodies. Cells were also labeled with APC-anti-Kit (clone 2B8, which recognizes an epitope distinct from the epitope detected by ACK2), eFluor 450 anti-Cd34, and PE-anti-Pdgfra. In experiments utilizing *Kit*$^{CreERT2/+}$*;R26*$^{mT-mG/mT-mG}$ mice, Cd34 was detected with Alexa Fluor (AF) 700-anti-Cd34 and Pdgfra was detected with Brilliant Violet 421-anti-Pdgfra. In additional aliquots of the same samples, fluorescent antibodies were replaced, one to three at a time, with isotype control antibodies conjugated with the corresponding fluorochrome. Samples were analyzed using a BD Biosciences (Becton, Dickinson and Company BD Biosciences, San Jose, CA) LSR II flow cytometer (see below for configuration) and FlowJo software (version 10; Tree Star, Ashland, OR).

**Antibodies used for flow cytometry analysis of cells freshly dissociated from murine gastric muscles**

| Target | Supplier | Host and clonality | Clone/ID | Isotype | Label | Final conc. or amount[a] / stomach |
|---|---|---|---|---|---|---|
| Cd16/32[b] | eBio | Rat mc[c] | 93 | $IgG_{2a}$, λ | | 1 µg |
| Cd11b[d] | eBio | Rat mc | M1/70 | $IgG_{2b}$, κ | PE[e] -Cy7[f] | 0.0312 µg |
| Cd45[g] | eBio | Rat mc | 30-F11 | $IgG_{2b}$, κ | PE-Cy7 | 0.0312 µg |
| F4/80[h] | eBio | Rat mc | BM8 | $IgG_{2a}$, κ | PE-Cy7 | 0.0625 µg |
| Kit[i] | eBio | Rat mc | ACK2 | $IgG_{2b}$, κ | APC[j] | 5 µg/mL |
| Kit | eBio | Rat mc | 2B8 | $IgG_{2b}$, κ | APC | 0.25 µg |
| Cd34[k] | eBio | Rat mc | RAM34 | $IgG_{2a}$, κ | AF[l] 700<br>or<br>eFluor 450 | 0.25 µg<br><br>0.2 µg |
| Pdgfra[m] | eBio | Rat mc | APA5 | $IgG_{2a}$, κ | BV[n] 421<br>or<br>PE | 0.5 µg<br><br>0.25 µg |

[a], Amount added to 100 µl staining volume; [b], Cd16, Fcgr3, Fc receptor, IgG, low affinity III; Cd32, Fcgr2b, Fc receptor, IgG, low affinity IIb; [c], mc, monoclonal; [d], Cd11b, Itgam, integrin alpha M; [e], PE, phycoerythrin; [f], Cy7, cyanine 7; [g], Cd45, Ptprc, protein tyrosine phosphatase, receptor type, C; [h], F4/80, Adgre1 (also known as Emr1), adhesion G protein-coupled receptor E1; [i], Kit, Kit oncogene; [j], APC, allophycocyanin; [k], Cd34, Cd34 antigen; [l], AF, Alexa Fluor;
[m], Pdgfra, platelet derived growth factor receptor, alpha polypeptide; [n], BV, Brilliant Violet. Supplier: eBio, Affymetrix eBioscience, Inc., San Diego, CA

**Configuration of the Becton Dickinson LSR II flow cytometer for cells freshly dissociated from murine gastric muscles**

| Laser | Excitation wavelength (nm) | Dichroic filter (nm) | Emission filter (nm; peak/bandwith) | Detector type | Use |
|---|---|---|---|---|---|
| Coherent® Sapphire™ 20 mW | 488 | | | Photodiode | Forward scatter |
| | | | 488/10 | PMT[a] | Side scatter |
| | | 505 LP[b] | 530/30 | PMT | mEGFP[c]; $DiBAC_4(3)$[d] |
| | | 550 LP | 575/26 | PMT | PE[e];<br>mTomato[f] |
| | | 595 LP | 610/20 | PMT | Unused |
| | | 685 LP | 695/40 | PMT | Trucount™ beads[g] |
| | | 735 LP | 780/60 | PMT | PE-Cy7[h] |
| Coherent® CUBE 100 mW | 407 | | 450/50 | PMT | eFluor 450;<br>BV[i] 421 |
| | | 505LP | 525/50 | PMT | Unused |
| | | 535 LP | 590/40 | PMT | Unused |
| | | 595 LP | 610/20 | PMT | Unused |
| | | 630 LP | 670/30 | PMT | Unused |
| | | 670 LP | 710/50 | PMT | Trucount™ beads |
| Coherent® CUBE 40 mW | 640 | | 660/20 | PMT | APC[j] |
| | | 685 LP | 712/20 | PMT | AF[k] 700 |
| | | 735 LP | 780/60 | PMT | Unused |

[a], PMT, photomultiplier tube; [b], LP, long-pass; [c], mEGFP, membrane-targeted enhanced green fluorescent protein; [d], $DiBAC_4(3)$, bis-(1,3-dibutylbarbituric acid)trimethine oxonol; [e], PE, phycoerythrin; [f], mTomato, membrane-targeted tandem dimer Tomato; [g], Trucount beads, multi-color fluorescent beads supplied as Trucount™ Absolute Counting Tubes, BD Biosciences, Becton, Dickinson and Company, San Jose, CA; [h], Cy7, cyanine 7; [I], BV, Brilliant Violet; [j], APC, allophycocyanin; [k], AF, Alexa Fluor

**Conditionally immortalized ICC.** ICL2A and ICL2B cells were isolated from gastric corpus+antrum *tunica muscularis* tissues of two homozygous, 12-day-old Immortomice by fluorescence-activated cell sorting (FACS) using a previously published and validated approach (*Approach 4* in ref.(Ordog et al., 2004b); see below for antibodies). Briefly, macrophages and ICC were labeled *in situ* by sequentially incubating the tissues with rat monoclonal anti-mouse F4/80 antibodies tagged with a tandem conjugate of PE and cyanine 5 (Cy5) (TRI-COLOR; 1 h at 37 ºC) and AF488-labeled ACK2 (3 h at 4 ºC), respectively. The tissues were dissociated by collagenase treatment and gentle trituration and passed through 30-μm polyester filters (Miltenyi Biotec, Auburn, CA). Macrophages, dendritic cells and other hematopoietic cells including $Kit^+$ mast cells were labeled with rat monoclonal anti-mouse/human Cd11b and hamster monoclonal anti-mouse Cd11c (Itgax) antibodies conjugated with superparamagnetic beads at 4 °C for 15 min, as well as with TRI-COLOR-conjugated rat monoclonal anti-mouse Cd11b and PE-Cy5-conjugated Cd45 antibodies for an additional 10 min at 4°C. The cells were then subjected to immunomagnetic selection (MACS; negative selection) using an MS magnetic column equipped with a 26-gauge needle as flow resistor and placed in a MiniMACS magnet (Miltenyi). Cells depleted of $Cd11b^+Cd11c^+$ cells were used for isolating ICC as $Kit^+Cd45^-Cd11b^-F4/80^-$ cells by FACS (**Figures 2A**). FACS was performed on a Beckman Coulter (Fullerton, CA) Elite sorter configured as described (Ordog et al., 2004b). The purified, $Kit^+Cd45^-Cd11b^-Cd11c^-F4/80^-$ ICC (56,000 cells) were plated at cloning density and cultured in SmGM-2 under conditions permissive for the tsTAg (33°C, 100 U/mL interferon-γ(Jat et al., 1991); Sigma-Aldrich). Two verified single-cell-derived clusters with ICC-like morphology were ring-cloned and propagated as ICL2A and ICL2B cells.

ICL1 cells were isolated by MACS as described previously (Ordog et al., 2004) with modifications: ICC were labeled *in situ* by incubating the gastric corpus and antrum *tunica muscularis* from a 116-day-old Immortomouse with monoclonal rat anti-mouse Kit antibody (clone: ACK2; 5 h at 4 ºC) conjugated with AF594. The tissues were dissociated by collagenase treatment and gentle trituration (Ordog et al., 2004; Ordog et al., 2004b). The resulting cell suspensions were reacted with goat $F(ab')_2$ anti-rat IgG conjugated with superparamagnetic beads, passed through 30-μm polyester filters and subjected to MACS (positive selection) twice using MS magnetic columns and Mini-MACS magnet (Miltenyi) (Ordog et al., 2004). The sorted cells were cultured and expanded in Clonetics Smooth Muscle Growth Medium-2 (SmGM-2; Lonza, Allendale, NJ) under conditions permissive for the tsTAg. A cluster dominated by cells with ICC-like morphology was ring-cloned and propagated (ICL1).

ICL1, ICL2A and ICL2B cells were maintained with SmGM-2 under permissive conditions (Jat et al., 1991). Cell proliferation was reduced or arrested by culturing with SmGM-2 under conditions semipermissive or nonpermissive for the tsTAg (39.5°C, 100 U/mL interferon-γ and 39.5°C, no interferon-γ, respectively) for 4 days (Bardsley et al., 2010; Jat et al., 1991). Reduction in tsTAg levels was verified by Western blotting and immunohistochemistry as described previously (Bardsley et al., 2010). These cell lines have not been authenticated or tested for Mycoplasma.

**Antibodies used for immunomagnetic and fluorescent sorting**

| Target | Supplier | Host and clonality | Clone/ID | Isotype | Label | Final conc. |
|---|---|---|---|---|---|---|
| Cd11b[a] | Miltenyi | Rat mc[b] | M1/70.15.11.5 | $IgG_{2b}$ | MB[c] | 10 µL/ $10^6$ cells |
| Cd11b | Thermo Fisher | Rat mc | M1/70.15 | $IgG_{2b}$ | TC[d] | 0.4 µg/ $10^6$ cells |
| Cd11c[e] | Miltenyi | Hamster mc | N418 | IgG | MB | 20 µL/ $10^6$ cells |
| Cd45[f] | eBio | Rat mc | 30-F11 | $IgG_{2b}$, κ | PE-Cy5[g] | 0.1 µg/ $10^6$ cells |
| F4/80[h] | Thermo Fisher | Rat mc | CI:A3–1 | $IgG_{2b}$ | TC | 4 µg/mL |
| Kit[i] | House | Rat mc | ACK2 | $IgG_{2b}$, κ | AF[j] 594 or AF 488 | 5 µg/mL |
| Rat IgG (H+L)[k] | Miltenyi | Goat pc | 130-048-501 | IgG $F(ab')_2$[l] | MB | 1:5 |

[a], Cd11b, Itgam, integrin alpha M; [b], mc, monoclonal antibody; [c], MB, superparamagnetic MicroBeads; [d], TC, TRI-COLOR, a tandem conjugate of phycoerythrin (PE) and cyanine 5 (Cy5); [e], Cd11c, Itgax, integrin alpha X; [f], Cd45, Ptprc, protein tyrosine phosphatase, receptor type, C; [g], PE-Cy5, generic PE-Cy5; [h], F4/80, Adgre1 (also known as Emr1), adhesion G protein-coupled receptor E1; [i], Kit, Kit oncogene; [j], AF, Alexa Fluor; [k], H+L, heavy and light chains; [l], $F(ab')_2$, $F(ab')_2$ fragment. Suppliers: Miltenyi, Miltenyi Biotec, Auburn, CA; Thermo Fisher, Thermo Fisher Scientific, Inc., Waltham, MA; eBio, Affymetrix eBioscience, Inc., San Diego, CA

**Immunocytochemistry and immunohistochemistry.** Cultured cells were fixed with 4% paraformaldehyde, permeabilized with Triton X-100 (Sigma-Aldrich) and immunolabeled as described previously(Bardsley et al., 2010) (see below for antibodies used). The permeabilization step was omitted from experiments designed to assess epitopes expressed on the cell surface (Kit, Cd34, Cd44, Cd45). For immunolabeling with the anti-Kit antibody ACK2, cells were fixed and permeabilized with acetone. Cells were examined using a Nikon (Melville, NY) Eclipse TS-100F fluorescent microscope equipped with Hoffman Modulation Contrast (Glen Cove, NY) objectives and a Jenoptik (Brighton, MI, USA) MFcool CCD digital camera. Specificity of immunolabeling was verified by omitting the primary antibodies, examining the samples with filter sets not designed for the fluorochrome used, preabsorption with the appropriate blocking peptide (Kcnn3, N-terminal and C terminal) and by Western blotting (Pdgfra, Kcnn3, C terminal and P2ry1). Post-acquisition modification of images was limited to maximum-transparency projection, assignment of pseudocolor, and adjustment of brightness and contrast, which were always applied to the entire image.

**Antibodies used in the immunocytochemistry and immunohistochemistry studies**

| Target | Supplier | Host and clonality | Clone/ID | Isotype | Label | Final conc. |
|---|---|---|---|---|---|---|
| Ano1[a] | Abcam | Rabbit pc[b] | Ab53212 | IgG | | 2 μg/mL |
| Cd34[c] | eBio | Rat mc[d] | RAM34 | $IgG_{2b}$,κ | | 5 μg/mL |
| Cd44[e] | eBio | Rat mc | IM7 | $IgG_{2b}$,κ | | 1.5 μg/mL |
| Cd45[f] | eBio | Rat mc | 30-F11 | $IgG_{2b}$,κ | | 2μg/mL |
| Gfap[g] | Sigma | Rabbit pc | G4546 | | | 1:100 |
| Kcnn3[h] (N-terminal) | Alomone | Rabbit pc | APC-025 | | | 1:500 |
| Kit[i] | House | Rat mc | ACK2 | $IgG_{2b}$,κ | | 5 μg/mL |
| Kit | eBio | Rat mc | 2B8 | $IgG_{2b}$,κ | | 5 μg/mL |
| Myh11[j] | BT | Rabbit pc | BT-562 | | | 1:100 |
| P2ry1[k] | Thermo Fisher | Rabbit pc | 34-7200 | IgG | | 0.1 μg/mL |
| Pdgfra[l] | Abcam, eBio | Rat mc | APA5 | $IgG_{2a}$ | | 1-2 μg/mL |
| Uchl1[m] | Cedarlane | Rabbit pc | CL95101 | IgG | | 1:1000 |
| Rat IgG (H+L)[n] | Thermo Fisher | Goat pc | A-11006 | IgG | AF[o] 488 | 5μg/mL |
| Rat IgG (H+L) | Thermo Fisher | Chicken pc | A-21471 | IgY | AF 594 | 5μg/mL |
| Rabbit IgG (H+L) | Thermo Fisher | Goat pc | A-11070 | IgG $F(ab')_2$[p] | AF 488 | 5μg/mL |
| Rabbit IgG (H+L) | Thermo Fisher | Chicken pc | A-21441 | IgY | AF 488 | 5μg/mL |
| Rabbit IgG (H+L) | Thermo Fisher | Chicken pc | A-21442 | IgY | AF 594 | 5μg/mL |
| Rabbit IgG | Millipore | Donkey pc | AP182F | IgG | FITC[q] | 1:200 |
| Rat IgG | Millipore | Donkey pc | AP189C | IgG | Cy3[r] | 1:400 |

[a], Ano1, anoctamin 1, calcium activated chloride channel; [b], pc, polyclonal antibody; [c], Cd34, Cd34 antigen; [d], mc, monoclonal antibody; [e], Cd44, Cd44 antigen; [f], Cd45, Ptprc, protein tyrosine phosphatase, receptor type, C; [g], Gfap, glial fibrillary acidic protein; [h], Kcnn3 (also known as SK3), potassium intermediate/small conductance calcium-activated channel, subfamily N, member 3; [i], Kit, Kit oncogene; [j], Myh11, myosin, heavy polypeptide 11, smooth muscle; [k], P2ry1, purinergic receptor P2Y, G-protein coupled 1; [l], Pdgfra, platelet derived growth factor receptor, alpha polypeptide; [m], Uchl1, ubiquitin carboxy-terminal hydrolase L1 (PGP9.5);
[n], H+L, heavy and light chains; [o], AF, Alexa Fluor; [p], $F(ab')_2$, $F(ab')_2$ fragment; [q], FITC, fluorescein isothiocyanate; [r], Cy3, cyanine 3. Suppliers: Abcam, Abcam plc, Cambridge, MA; eBio, Affymetrix eBioscience, Inc., San Diego, CA; Sigma, Sigma-Aldrich Corp., St. Louis, MO; Alomone, Alomone Labs, Ltd., Jerusalem, Israel; BT, Biomedical Technologies, Alfa Aesar, Ward Hill, MA; Thermo Fisher, Thermo Fisher Scientific, Inc., Waltham, MA; Cedarlane, Burlington, NC; Millipore, EMD Millipore Corp., Billerica, MA

**Primers used in reverse transcription–polymerase chain reaction (RT-PCR) studies (Organism: mus musculus)**

| Gene symbol | Description | Primer sequences (forward/reverse) |
|---|---|---|
| *Kit*[a] | Kit oncogene | CGCCTGCCGAAATGTATG |
| | | GGTTCTCTGGGTTGGGGT |
| *Kit*[b] | Kit oncogene | AGGCTATCCCTGTTGTGTCTGTGC |
| | | GCTTTACCTGGGCTATGTGCTGAG |
| *Ano1* | Anoctamin 1, calcium activated chloride channel | GGGAACCCCCATGGTCAGAACACA |
| | | TCTGCTGGCTGATGTCTTTGGGGA |
| *Cacna1c* | Calcium channel, voltage-dependent, L type, alpha 1C subunit | CACCCCATTATCCAGCCTCG |
| | | TCAATGGAGCGGTACAGCAG |
| *Cacna1d* | Calcium channel, voltage-dependent, L type, alpha 1D subunit | ACACCACGATTGCCCTACAG |
| | | TTCCAAGCAGGGCACCATTT |
| *Cacna1h* | Calcium channel, voltage-dependent, T type, alpha 1H subunit | CCCTGGATGCTGCTCTACTTC |
| | | CGTGTGTGAATAGTCTGCGT |
| *Cav1* | Caveolin 1, caveolae protein | AGGTGACTGAGAAGCAAGTGT |
| | | AACTGTGTGTCCCTTCTGGTT |
| *Trpm7* | Transient receptor potential cation channel, subfamily M, member 7 | TTCTACTTCTCTTCACTCGGTGC |
| | | TTCTGGAGAGTCTTCCGTCG |
| *Slc8a1* | Solute carrier family 8 (sodium/calcium exchanger), member 1 | GAGATTGGAGAGCCCCGTCT |
| | | TCAGTGGCTGCTTGTCATCAT |
| *Gucy1a3* | Guanylate cyclase 1, soluble, alpha 3 | ATGTCAGCCCCACCACATAC |
| | | CCCTGACGCTTTGCCTAAGA |
| *Gucy1b3* | Guanylate cyclase 1, soluble, beta 3 | CACTGAGAGCCTTGGAGGATG |
| | | TGTCGTATCTTTTGGCAGGCA |
| *Prkcq* | Protein kinase C, theta | TTGAAAGCACCCAACAGGCT |
| | | GCTCACTTCGGGTTCAGGAG |
| *Atp2b2* | ATPase, Ca++ transporting, plasma membrane 2 | TCCATCAATGCCAAGACGCT |
| | | CTCTGTCTTGTTGCCCACCT |
| *Actb* | Actin, beta | ATGGTGGGAATGGGTCAGAAGG |
| | | GCTCATTGTAGAAGGTGTGGTGCC |
| *Chrm2* | cholinergic receptor, muscarinic 2, cardiac | GCCAGACTCCACCAGATCG |
| | | TGTGTTCAGTAGTCAAGTGGC |
| *Chrm3* | cholinergic receptor, muscarinic 3, cardiac | CTCCTCTTGAAGTGCTGCGT |
| | | GTTGGGAAACAAAGGCGAGG |
| *P2ry6* | pyrimidinergic receptor P2Y, G-protein coupled | GCTGTGTCAGAGGGAGTTTT |
| | | CCAGATTTGGTGTTCCTCGT |
| *Etv1 v1* | ets variant 1, isoform a | CTTCCAAGGGGAAGTGCTGGGC |
| | | CTCGGTACAGTTTCTCCCACGCT |
| *Etv1 v2* | ets variant 1, isoforms a and b | AAGTGCAGGCGTCTTCTTCCCTCC |
| | | TGGAGTGCTGGATGGTGTCGGG |
| *Trpc2* | transient receptor potential cation channel, subfamily C, member 2 | GCCCTACCAGGAATCCGAGA |
| | | TGCTCTTCCATGCCGAACAT |
| *Trpc1* | transient receptor potential cation channel, subfamily C, member 1 | AGCCTCTTGACAAACGAGGA |
| | | TACGGCGGTAACCTGACATC |
| *Acvr2a* | activin receptor IIA | TCCAGCCAACAACCTTGCTTCAC |
| | | GCGCCTCGGGAAAATGGGAGC |
| *Lrig1* | leucine-rich repeats and immunoglobulin-like domains 1 | CCCATCCAGAGTCGCTGAAG |
| | | CAGGCGAAGGTCATTGGTGA |
| *Cav2* | caveolin 2 | TTGCGGGTATCCTGTTTGCTA |
| | | GAAGGCAAGACCATTAGGCAG |
| *Prkg1* | protein kinase, cGMP-dependent, type I | GGAGTTGGAGGTTTCGGACG |

| | | |
|---|---|---|
| | | GATGTGCTCCTGCTGTCTGG |
| *Atp2a2* | ATPase, Ca++ transporting, cardiac muscle, slow twitch 2 | AGGCACACTTACCACAAACCA |
| | | GAGCACAGATGGTGGCTAACT |
| *Itpr1* | inositol 1,4,5-trisphosphate receptor 1 | AGCATCCAGAGACTACCGAA |
| | | AACATCCACGAGCACAGACAG |
| *Pdgfra* | Platelet derived growth factor receptor, alpha polypeptide | TCGCCAAAGTGGAAGAGACC |
| | | CACCAGGACGATGAGAGAGA |
| *Kcnn3* | potassium intermediate/small conductance calcium-activated channel, subfamily N, member 3 | AAGCCAACACCCTGGTGGACCT |
| | | AGGGGCGTGGCTAGTTCCCA |
| *P2ry1* | purinergic receptor P2Y, G-protein coupled 1 | TGAAGGCACGAGATCCTAGC |
| | | GCACCTACGGCATCAACAAA |

[a], Exons 19-21 (intracellular; used in qualitative RT-PCR experiments); [b], Exons 4-5 (extracellular)

**Antibodies used for flow cytometry analysis of conditionally immortalized cells**

| **Target** | **Supplier** | **Host and clonality** | **Clone** | **Isotype** | **Label** | **Amount for 4x10^6 cells** |
|---|---|---|---|---|---|---|
| Cd16/32[a] | eBio | Rat mc[b] | 93 | $IgG_{2a}$, λ | | 1 μg |
| Cd34[c] | eBio | Rat mc | RAM34 | $IgG_{2a}$, κ | PE[d] | 0.2 μg |
| Pdgfra[e] | eBio | Rat mc | APA5 | $IgG_{2a}$, κ | APC[f] | 0.125 μg |
| Kit[g] | eBio | Rat mc | ACK2 | $IgG_{2b}$, κ | APC-Cy7[h] | 0.25 ug |
| Kit | eBio | Rat mc | 2B8 | $IgG_{2b}$, κ | APC-Cy7 | 0.25 μg |
| Cd44[i] | BioLegend | Rat mc | IM7 | $IgG_{2b}$, κ | APC-Cy7 | 0.0625 μg |
| Kit | eBio | Rat mc | ACK2 | $IgG_{2b}$, κ | PE-Cy7 | 2.0 ug |
| Kit | eBio | Rat mc | 2B8 | $IgG_{2b}$, κ | PE-Cy7 | 2.0 ug |

[a], Cd16, Fcgr3, Fc receptor, IgG, low affinity III; Cd32, Fcgr2b, Fc receptor, IgG, low affinity IIb; [b], mc, monoclonal; [c], Cd34 antigen; [d], PE, phycoerythrin; [e], Pdgfra, platelet derived growth factor receptor, alpha polypeptide; [f], APC, allophycocyanin; [g], Kit, Kit oncogene; [h], Cy7, cyanine 7; [i], Cd44, Cd44 antigen. Suppliers: eBio, Affymetrix eBioscience, Inc., San Diego, CA; BioLegend, San Diego, CA

**Antibodies used in Western immunoblotting studies**

| Target | Supplier | Host and clonality | Clone/ID | Isotype | Label | Final conc. |
|---|---|---|---|---|---|---|
| Gapdh[a] | Novus | Goat pc[b] | IMG-3073 | | | 0.05 μg/mL |
| Gapdh | Sigma | Rabbit pc | G9545 | | | 1:40000 |
| SV40[c] | BD | Mouse mc[d] | PAb101 | $IgG_{2a}$ | | 0.2 μg/mL |
| Kit[e] | RDS | Goat pc | AF1356 | IgG | | 0.2 μg/mL |
| Pdgfra[f] | CST | Rabbit pc | #3164 | | | 1:2000 |
| P2ry1[g] | Thermo Fisher | Rabbit pc | 34-7200 | IgG | | 0.5 μg/mL |
| Kcnn3[h] (C-terminal) | Alomone | Rabbit pc | APC-103 | | | 1:1000 |
| Myh11[i] | BT | Rabbit pc | BT-562 | | | 1:15000 |
| Uchl1[j] | Cedarlane | Rabbit pc | CL95101 | IgG | | 1:15000 |
| Ezh2[k] | CST | Rabbit pc | #4905 | | | 1:2000 |
| H3K27me3[l] | CST | Rabbit mc | C36B11 | IgG | | 1:10000 |
| H3 | Thermo Fisher | Mouse mc | 865R2 | IgM, κ | | 0.5 μg/mL |
| Suz12[m] | SCBT | Goat pc | sc-46264 | IgG | | 1:1000 |
| Rabbit IgG (H+L)[n] | LI-COR | Donkey pc | 926-32223 | IgG | IRDye 680 | 1:10000 |
| Mouse IgG (H+L) | LI-COR | Donkey pc | 926-32222 | IgG | IRDye 680 | 1:10000 |
| Rabbit IgG (H+L) | LI-COR | Donkey pc | 926-32213 | IgG | IRDye 800CW | 1:10000 |
| Goat IgG (H+L) | LI-COR | Donkey pc | 926-32214 | IgG | IRDye 800CW | 1:10000 |

[a], Gapdh, glyceraldehyde-3-phosphate dehydrogenase; [b], pc, polyclonal antibody; [c], SV40, simian virus 40 large T antigen; [d], mc, monoclonal antibody; [e], Kit, Kit oncogene; [f], Pdgfra, platelet derived growth factor receptor, alpha polypeptide; [g], P2ry1, purinergic receptor P2Y, G-protein coupled 1; [h], Kcnn3 (also known as SK3), potassium intermediate/small conductance calcium-activated channel, subfamily N, member 3; [i], Myh11, myosin, heavy polypeptide 11, smooth muscle; [j], Uchl1, ubiquitin carboxy-terminal hydrolase L1 (PGP9.5); [k], Ezh2, enhancer of zeste homolog 2 (Drosophila); [l], H3K27me3, histone H3 lysine 27, trimethylated; [m], Suz12, suppressor of zeste 12 homolog (Drosophila); [n], H+L, heavy and light chains. Suppliers: Novus, Novus Biologicals, Littleton, CO; Sigma, Sigma-Aldrich Corp., St. Louis, MO; BD, BD Biosciences, Becton, Dickinson and Company, San Jose, CA; RDS, R&D Systems, Inc., Minneapolis, MN; CST, Cell Signaling Technology, Inc., Beverly, MA; Thermo Fisher, Thermo Fisher Scientific, Inc., Waltham, MA; Alomone, Alomone Labs, Ltd., Jerusalem, Israel; BT, Biomedical Technologies, Alfa Aesar, Ward Hill, MA; Cedarlane, Burlington, NC; SCBT, Santa Cruz Biotechnology, Inc., Dallas, TX; LI-COR Biosciences, Lincoln, NE

**Antibodies used in ChIP and ChIP-seq studies**

| Target | Supplier | Host and clonality | Clone/ID | Isotype | Amount for 1x10^6 cells |
|---|---|---|---|---|---|
| NST[a] | Millipore | Rabbit pc[b] | 12-370 | IgG | 2 µg |
| H3K27me3[c] | CST | Rabbit mc[d] | C36B11/#9733 | IgG | 2 µg |
| H3K27ac[e] | Abcam | Rabbit pc | ab4729 | IgG | 2 µg |
| H3K4me3[f] | Abcam | Rabbit pc | ab8580 | IgG | 2 µg |
| H3K9me3[g] | Abcam | Rabbit pc | ab8898 | IgG | 2 µg |
| Ezh2[h] | CST | Rabbit mc | D2C9/#5246 | IgG | 2 µg |

[a], NST, no specific target; [b], pc, polyclonal; [c], H3K27me3, histone H3 lysine 27, trimethylated; [d], mc, monoclonal; [e], H3K27ac, histone H3 lysine 27, acetylated; [f], H3K4me3, histone H3 lysine 4, trimethylated; [g], H3K9me3, histone H3 lysine 9, trimethylated; [h], Ezh2, enhancer of zeste homolog 2 (Drosophila). Suppliers: Millipore, EMD Millipore Corp., Billerica, MA; CST, Cell Signaling Technology, Inc., Beverly, MA; Abcam, Abcam plc, Cambridge, MA

**Primers targeting transcription start sites used in ChIP-qPCR studies**

| Organism | Gene symbol | Description | Primer sequences (forward/reverse) |
|---|---|---|---|
| *Mus musculus* | *T* | Brachyury | GAGACGCCGATCCGCCGAAG |
| | | | ACTCTCCACTCCCACGCGCT |
| *Mus musculus* | *Actb* | Actin, beta | CCCGGCAAGCCGAATAGGCA |
| | | | ACCAGACGCTACGATCACGCC |
| *Mus musculus* | *Kit* | Kit oncogene | CACTTGGGCGAGAGCTGTA |
| | | | GAGGGTGCAGTCCTCTTGTC |
| *Mus musculus* | *Ano1* | Anoctamin 1, calcium activated chloride channel | CCCAACTCTAGCTTGGCCC |
| | | | CCACAAACTGGCAGAGTGGT |
| *Mus musculus* | *Cacna1h* | Calcium channel, voltage-dependent, T type, alpha 1H subunit | CTTGGTCCGTGTTTCTCCGT |
| | | | CTCTCCCGCGAGACAGTATG |
